# Extrafollicular B cell responses are associated with autoreactivity against the slit diaphragm in idiopathic podocytopathies

**DOI:** 10.1101/2025.09.06.674608

**Authors:** Tho-Alfakar Al-Aubodah, Simon Leclerc, Lamine Aoudjit, Zhiyang Liu, Ratna Samanta, Cherry Mammen, Allison Dart, Anne-Laure Lapeyraque, Catherine Morgan, Mallory L. Downie, Kunimasa Yan, Satoka Mori, Susan M. Samuel, Ciriaco A. Piccirillo, Tomoko Takano

## Abstract

Idiopathic podocytopathies lack mechanistic biomarkers and rely on empirical immunosuppression, yet the recent discovery of autoantibodies against slit diaphragm proteins implicates podocyte-directed autoimmunity as a driver of podocyte injury. However, the immune mechanisms underlying autoreactivity and phenotypic heterogeneity, particularly across age groups, remain unclear. Here, we identify expansion of CD21^low^ T-bet^+^ atypical B cells (atBCs) in peripheral blood as a shared immunologic feature across idiopathic podocytopathies. In children, atBC accumulation was associated with heightened B cell activation, favoring extrafollicular responses and plasmablast differentiation. Serologic profiling revealed that autoantibodies against the slit diaphragm protein CRB2 are highly prevalent across disease phenotypes and frequently co-occur with anti-Nephrin antibodies, particularly in adults. Autoantibody abundance correlated with atBC expansion in affected children, implicating extrafollicular B cell responses as a key pathway for their generation, whereas adults showed additional involvement of follicular pathways. Complementary mouse modeling demonstrated intermolecular epitope spreading as a mechanism linking anti-CRB2 and anti-Nephrin autoreactivity. Together, these findings define a shared immune framework for idiopathic podocytopathies in which extrafollicular B cell responses drive autoreactivity to multiple slit diaphragm proteins, with implications for biomarker development and therapeutic stratification.

## INTRODUCTION

Idiopathic podocytopathies are kidney diseases characterized by injury to podocytes, the specialized epithelial cells that form the glomerular filtration barrier, resulting in heavy proteinuria and nephrotic syndrome (*1*). These diseases encompass a spectrum of clinical and histologic phenotypes that, in the absence of a defined etiology, are currently classified by treatment response and biopsy findings rather than by underlying pathogenesis (*2, 3*). In children, where kidney biopsies are infrequently performed, disease is typically diagnosed clinically as idiopathic nephrotic syndrome (INS) and subdivided into steroid-sensitive (SSNS, ∼90% of cases) and steroid-resistant (SRNS) forms. When biopsy is done, most children and a minority of adults show minimal change disease (MCD), whereas most adults exhibit more severe podocyte injury with glomerular sclerosis, leading to a diagnosis of focal segmental glomerulosclerosis (FSGS), which often responds poorly to corticosteroids. Despite this heterogeneity, idiopathic podocytopathies likely represent a disease continuum, as MCD can progress to FSGS and recurrent FSGS after transplantation initially manifests with MCD-like lesions (*4*).

Recent identification of autoantibodies targeting Nephrin – a major component of the podocyte slit diaphragm – has repositioned idiopathic podocytopathies as autoimmune diseases characterized by podocyte-directed humoral immunity (*5, 6*). Anti-Nephrin autoantibodies are highly prevalent in children at disease onset but show variable seroprevalence in relapses and in adults, suggesting that additional immune mechanisms contribute to disease activity and progression (*6*). Consistent with this notion, autoantibodies against two other slit diaphragm proteins, namely Podocin and Kirrel1, have also been described, suggesting slit diaphragm-directed autoimmunity as a central feature of idiopathic podocytopathies (*7, 8*). This contrasts with membranous nephropathy, in which humoral responses predominantly target subepithelial podocyte antigens such as PLA2R1, THSD7A, and NELL1 (*9–11*). However, the scope of slit diaphragm autoreactivity and the immune pathways that generate responses across idiopathic podocytopathies remain poorly defined. We recently demonstrated that immunization of mice with Crumbs cell polarity complex 2 (CRB2), a slit diaphragm protein essential for Nephrin localization and filtration barrier integrity, induces anti-CRB2 autoantibodies, podocyte injury, and proteinuria (*12*). Although CRB2 autoreactivity has not been established in humans, pathogenic variants in CRB2 cause congenital FSGS, underscoring its relevance to podocyte biology and disease (*13, 14*). Whether CRB2 represents an additional target of humoral autoimmunity in idiopathic podocytopathies is unknown.

The clinical efficacy of B cell–targeted therapies further support a pathogenic role for humoral immunity in these diseases. CD20-based B cell depletion with rituximab or ofatumumab is effective in many patients with INS, MCD, and some forms of FSGS, particularly in children, although responses are heterogeneous and often diminished in adults (*7–9*). Despite this, the nature of the podocytopathic B cell response remains uncharacterized. Prior immunophenotyping studies described elevated levels of CD27^+^ memory B cells (MBCs) and plasmablasts in circulation, the latter primarily in children and adults prior to treatment, indicative of a dysregulated B cell compartment (*10–14*). We recently showed that children with INS have an enlarged pool of CD21^low^ T-bet^+^ atypical B cells (atBCs) and marginal zone (MZ)-like B cells in peripheral blood, subsets that are often enriched with autoreactive clones and are frequently associated with systemic autoimmunity (*14, 15*). CD21^low^ T-bet^+^ atBCs, in particular, contribute to autoantibody production in systemic lupus erythematosus and arise primarily through extrafollicular B cell responses (*16–18*). Unlike germinal centers that yield isotype-switched MBCs with highly affined BCRs and long-lived plasma cells, extrafollicular responses are faster, with strong BCR signals promoting early plasmablast formation to provide a rapid source of IgM and IgG antibody (*19*). Notably, while long-lived plasmablasts tend to exhibit greater steroid resistance and are completely insensitive to CD20-based depletion strategies, short-lived plasmablasts are steroid-sensitive and remain susceptible to CD20-based depletion as they rely on renewal from their MBC precursors (*20, 21*). Whether extrafollicular responses play a role in podocyte-directed autoimmunity remains unknown.

Here, we comprehensively profile peripheral B cell states and podocyte-directed autoantibody responses in children and adults with idiopathic podocytopathies, revealing a shared immunologic architecture across disease phenotypes. We identify expansion of CD21^low^ T-bet^+^ atypical B cells and plasmablasts as a conserved feature of disease, and demonstrate that in INS, B cells are transcriptionally primed for extrafollicular activation, characterized by heightened BCR signaling and enrichment of VH4-39^+^ clones associated with autoreactivity and polyreactivity. Extending beyond Nephrin, we show that autoantibodies targeting the slit diaphragm protein CRB2 are particularly prevalent in adult disease and frequently co-occur with anti-Nephrin responses, indicating a broader slit diaphragm-directed humoral repertoire than previously appreciated. Importantly, the abundance of anti-CRB2 and anti-Nephrin autoantibodies correlates with expansion of CD21^low^ T-bet^+^ atypical B cells and isotype-switched plasmablasts in children with INS, thus linking extrafollicular subsets to autoreactivity. Together, these findings implicate extrafollicular B cell responses as a central mechanism driving podocyte-directed autoimmunity and provide a mechanistic framework for the clinical efficacy of B cell-targeted therapies, while identifying immunologic and serologic features that may inform future biomarker development and therapeutic stratification.

## RESULTS

### CD21^low^ T-bet^+^ B cells are expanded in idiopathic podocytopathies

We first aimed to characterize the B cell landscape in children and adults with idiopathic podocytopathies. To this end, we obtained blood samples from 26 children and 17 adults during active disease and compared the composition of their B cell compartments to age-matched healthy controls (**table S1, fig. 1A, fig. S1A**). Only one individual analyzed (NS26, steroid-resistant FSGS) had previously received rituximab, though they had successfully reconstituted their B cell compartment and continued to relapse by time of sampling (**table S1**). Total B cell frequencies were comparable to those in healthy individuals (**fig. S1B**). Nevertheless, naïve B cell frequencies were greatly diminished in affected children and adults, though at distinct stages: children exhibited a selective decrease in pre-splenic transitional B cells, whereas adults had fewer post-splenic mature B cells (**fig. 1B, C, fig. S1B**). Comparable frequencies of classical (CD27^+^ CD21^+^) MBCs were present in affected and healthy children, whereas isotype-switched MBCs, indicating a germinal center origin, were notably expanded in adults (**fig. 1D**).

**Figure 1.**
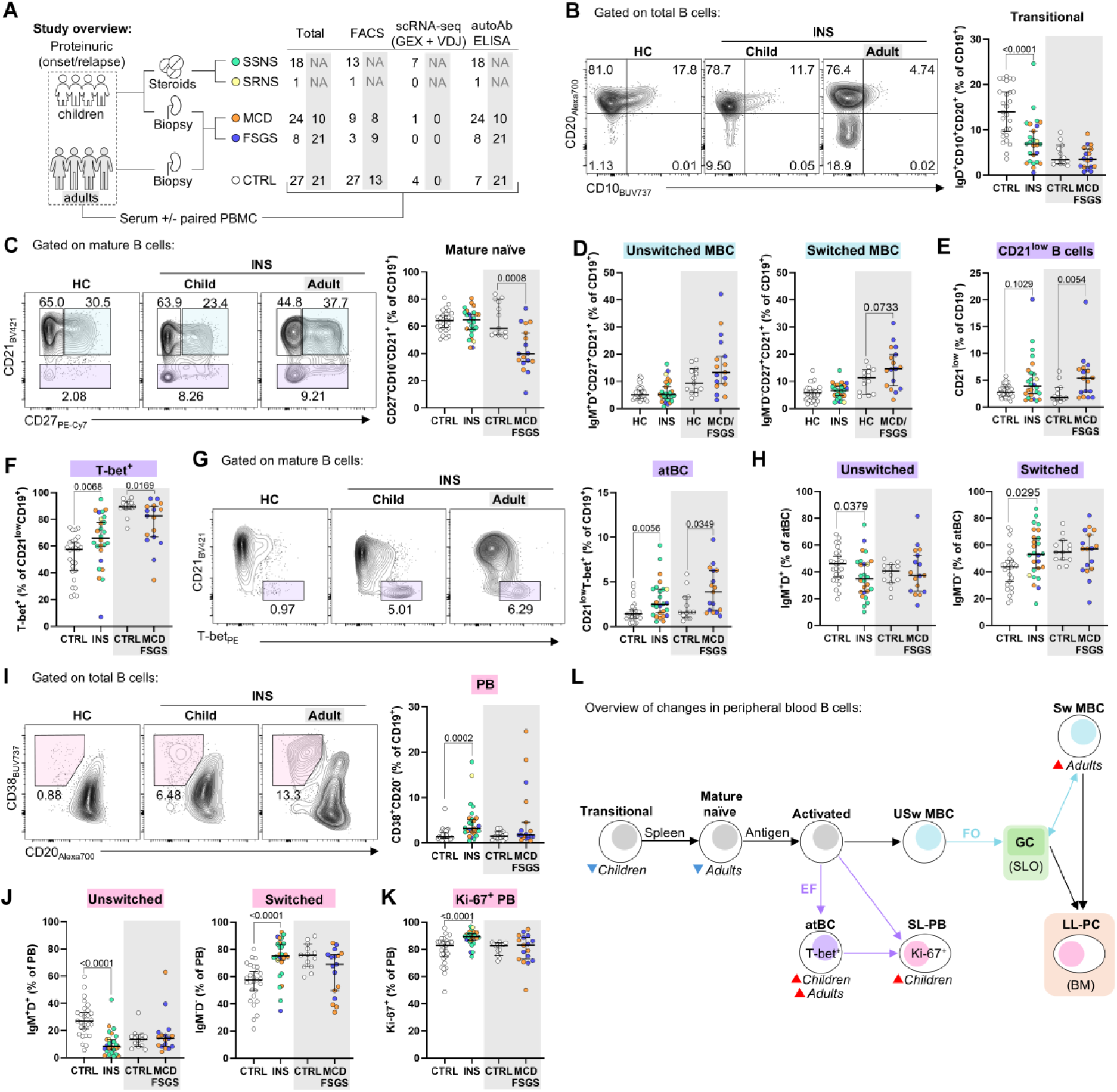
T-bet^+^ CD21^low^ atypical B cells and plasmablasts are expanded in children and adults with idiopathic podocytopathies. (**A**) Overview of the study with adult data highlighted in gray. SSNS, steroid-sensitive nephrotic syndrome; SRNS, steroid-resistant nephrotic syndrome; MCD, minimal change disease; FSGS, focal segmental glomerulosclerosis; FACS, fluorescence-activated cell sorting; scRNA-seq, single-cell RNA-sequencing; GEX, gene expression; VDJ, variable-diversity-joining; autoAb, autoantibody. (**B**) Representative flow plots and quantification of frequencies of transitional naïve B cells (IgD^+^ CD10^+^ CD20^+^) within total B cell pool. (**C**) Representative flow plots of mature B cells (CD10^−^ CD20^+^) highlighting naïve (CD21^+^ CD27^−^), memory (MBC, CD21^+^ CD27^+^), and CD21^low^ compartments alongside frequencies of mature naïve B cells within total B cell pool. (**D-E**) Frequencies of unswitched (D), isotype-switched (D), and CD21^low^ B cells (E) within total B cell pool. (**F**) Frequencies of CD21^low^ B cells expressing T-bet. (**G**) Representative flow plots and quantification of frequencies of CD21^low^ T-bet^+^ atypical B cells (atBCs). (**H**) Percentage of CD21^low^ T-bet^+^ atBCs that are unswitched or isotype-switched. (**I**) Representative flow plots and quantification of frequencies of plasmablasts (PB, CD38^+^ CD27^+^). (**J**) Percentage of plasmablasts that are unswitched or isotype-switched. (**K**) Percentage of plasmablasts that are actively cycling. (**L**) Diagram of peripheral B cell development outlining the major changes observed. Each point refers to a single donor and summary statistics are expressed median with 95% confidence intervals. *P*-values were determined using Mann-Whitney *U*-tests comparing controls and patients in children and adults separately.

CD21^low^ B cells were modestly expanded in children with INS, as we previously showed (*14*), and were prominent in adults with MCD/FSGS (**fig. 1E**). This compartment showed higher frequencies of T-bet^+^ B cells (atBCs) in children with INS than healthy children, whereas CD21^low^ B cells in both adults with MCD/FSGS and healthy adults were nearly entirely T-bet^+^ (**fig. 1F**). Indeed, frequencies of CD21^low^ T-bet^+^ atBCs were significantly elevated in both affected children and adults, establishing their expansion as a common feature of idiopathic podocytopathies (**fig. 1G, fig. S1B**). In childhood INS, atBCs showed greater rates of isotype-switching than in health, reaching levels observed in adults (**fig. 1H**). Double-negative (DN, IgD^−^ CD27^−^), particularly DN2 (CXCR5^low^ CD21^low^), B cells are a partially overlapping population with CD21^low^ T-bet^+^ atBCs that are expanded in systemic lupus erythematosus (*16*). These subsets were not significantly enriched in idiopathic podocytopathies (**fig. S1C**). Instead, many CD21^low^ T-bet^+^ atBCs in idiopathic podocytopathies retained surface IgD. Consistently, the DN2 precursor population, termed activated naïve (aNAV, IgD^+^ CD27^−^ CXCR5^low^ CD21^low^), which has surface IgD expression was significantly expanded in adults with MCD/FSGS (**fig. S1C**).

Plasmablasts were also significantly expanded in affected children (**fig. 1I, fig. S1B**). Plasmablasts in INS had higher rates of isotype-switching, comparable to levels observed in adults, and greater cycling (**fig. 1J, K**). Only a few adults with MCD/FSGS exhibited plasmablast expansion (**fig. 1I**). Together, these findings point to a dysregulated peripheral B cell landscape in idiopathic podocytopathies characterized by a contraction of the naïve compartment and the expansion of atBCs, with adults further showing evidence of germinal center involvement (**fig. 1L**).

### B cells in INS are primed for extrafollicular activation and conversion into plasmablasts

While CD21^low^ T-bet^+^ atBC expansion was previously linked to extrafollicular responses in systemic autoimmunity, the underlying factors promoting their expansion in idiopathic podocytopathies remain uncharacterized. To this end, we conducted single-cell gene expression and V(D)J repertoire profiling on circulating mature (CD10-depleted) B cells from children with INS, considering this cohort to be at an earlier stage of B cell dysregulation (**table S1**). B cells from three age-matched healthy children and one child with a non-glomerular atopic disease were profiled in parallel as healthy and disease controls, respectively. We obtained 118,706 high quality B cells (93,766 from INS; 24,310 from controls) of which 88,334 cells (69,843 from INS; 18,491 from controls) had paired V(D)J sequences (**fig. S2A, B**). Integrated clustering yielded 11 distinct B cell populations: five naïve (B0, B1, B3, B4, B9) defined by high *BACH2* and *IL4R* expression, five memory (B2, B5, B6, B7, B8) that expressed memory-associated transcripts like *BCL2A1*, *TNFRSF13B*, and *IGHA1*, and a single cluster (B10) of plasmablasts expressing *PRDM1*, *JCHAIN*, and *XBP1* (**fig. 2A, B, fig. S2C, data file S1**). Notably, the naïve B3 and memory B7 clusters had prominent expression of activation genes (e.g. *FOSB*, *NR4A1*, *CD69*) and were thus accordingly termed actNaïve and actMBC (**fig. 2B, fig. S2C**).

**Figure 2.**
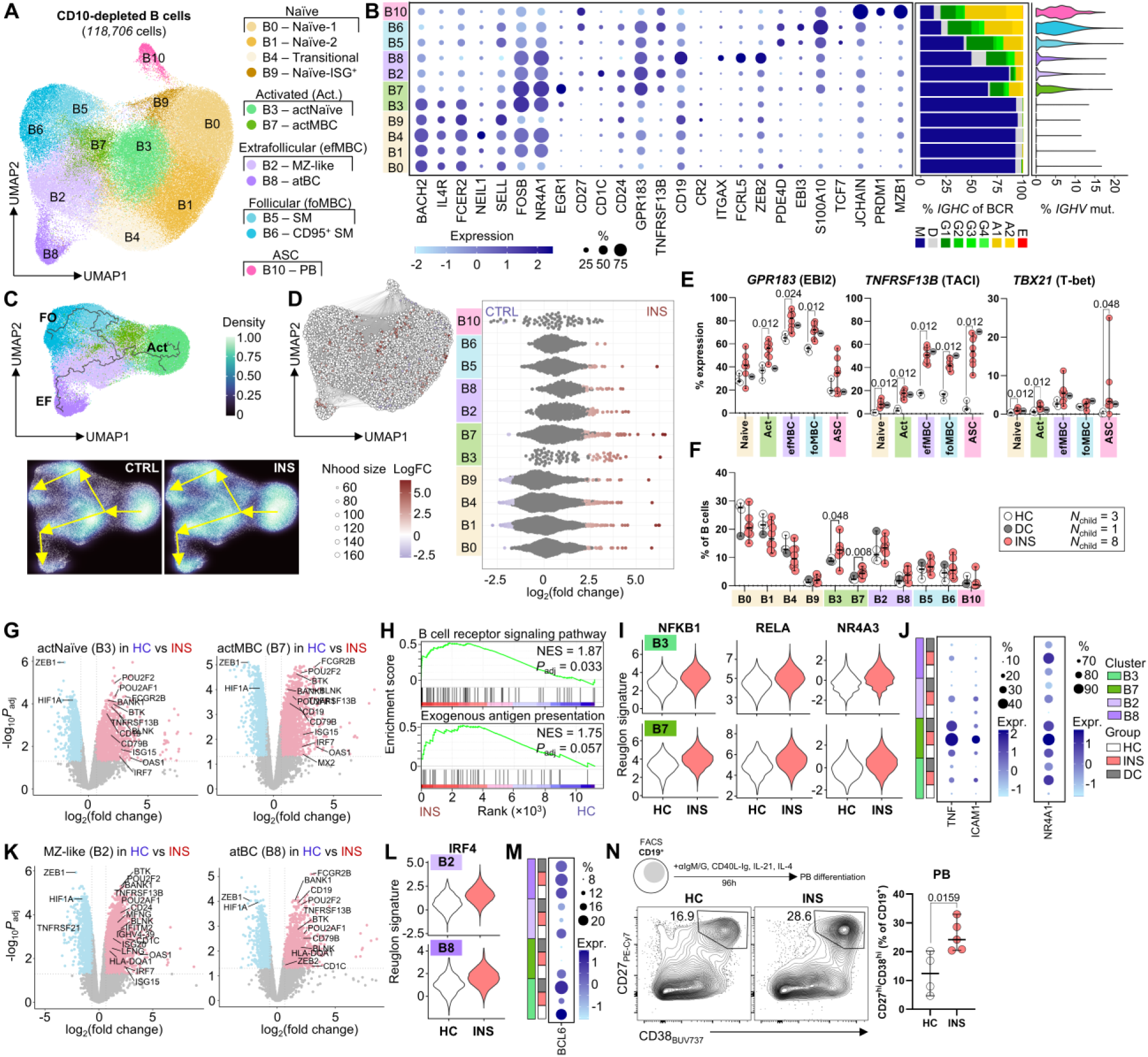
B cell hyperactivity in INS favours extrafollicular activation and plasmablast formation. (**A**) Uniform manifold approximation and projection (UMAP) plot of B cells from children with INS (*N* = 8), healthy controls (HC, *N* = 3), and a disease control (DC, *N* = 1). **(B)** Bubble plot showing expression intensity and frequency of key genes relating to B cell cluster identities. *IGHC* gene usage by each cluster are expressed as percentages in the stacked bar graph, and somatic hypermutation is shown using a violin plot depicting percentages of mutated nucleotides in the *IGHV* gene. (**C**) UMAP of activated and MBCs showing trajectories inferred using Monocle3. Density plots of B cells from controls (CTRL) and children with INS are shown below with yellow arrows showing the distinct trajectories. EF, extrafollicular; FO, follicular. (**D**) Distinct states, or neighbourhoods, of B cells are plotted on the UMAP of total B cells and those that are significantly (spatial *FDR* < 0.1) differentially abundant between INS and control children are coloured by their log_2_(fold change). Quantification for each cluster is shown. **(E)** Percentage of cells in each cluster expressing *EBI2*, *TNFRSF13B*, and *TBX21*. (**F**) Frequencies of B cells within each cluster. (**G**) Volcano plots of differentially expressed genes between INS and healthy controls within actNaïve and actMBC clusters. (**H**) Enrichment of gene ontology pathways in activated B cells from INS versus healthy controls. (**I**) Violin plots depicting the collecTRI NFKB1, RELA, and NR4A3 regulon signatures in actNaïve and actMBC clusters in INS and healthy controls. (**J**) Bubble plots showing the expression intensity and frequency of NFκB target genes *TNF* and *ICAM*, as well as *NR4A1*. (**K**) Volcano plots of differentially expressed genes between INS and healthy controls within MZ-like B cell and atBC clusters. (**L**) Violin plots depicting the collecTRI IRF regulon signature in MZ-like B cells and atBCs in INS and healthy controls. (**M**) Bubble plots showing the expression intensity and frequency of *BCL6*. (**N**) Schematic of the *in vitro* B cell stimulation with representative flow plots showing plasmablast formation and quantification. In E, F, and N, each dot represents a single donor and summary statistics are represented as median with 95% confidence intervals. *P*-values were determined using independent Mann-Whitney *U*-tests between healthy controls and children with INS. In D, comparisons were done between INS and all controls.

Memory clusters were further divided into extrafollicular memory B cells (efMBC: B2, B8) defined by low class-switch recombination and low somatic hypermutation, and germinal center-derived/follicular MBCs (foMBC: B5, B6) with high isotype-switching and mutation rates. Cluster B2 resembled MZ B cells defined by high expression of *CD1C*, *TNFRSF13B*, *CD24*, and IgM BCRs (**fig. 2B**), while cluster B8 corresponded with CD21^low^ atBCs given their highly characteristic transcriptional profile (*FCRL5*, *ITGAX*, *ZEB2*, etc.) (**fig. 2B, fig. S2C**) (*14, 22–25*). Both B5 and B6 corresponded with isotype-switched memory (SM) B cells, with B6 (CD95^+^ SM) corresponding to the recently described *FAS*-expressing MBCs that display the highest rates of isotype-switching and somatic hypermutation (**fig. 2B**). Trajectory analysis of activated and memory B cells showed distinct pathways along efMBC and foMBC clusters, corresponding with extrafollicular and germinal center-dependent activation routes (**fig. 2C, D**). B cells in INS showed increased bias for the extrafollicular pathway, and this was confirmed by differential abundance testing where efMBC subsets encompassed more INS-associated states than foMBC subsets (**fig. 2D**). In line with this, expression of genes encoding EBI2 (*GPR183)*, a homing receptor that retains B cells within extrafollicular niches, and the BAFF/APRIL receptor TACI *(TNFRSF13B*), both of which are essential for extrafollicular responses, was more prevalent in B cells from INS than healthy controls (**fig. 2E**) (*19, 26*). This increased expression was notably also observed in naïve and activated B cells suggesting a bias for extrafollicular activation prior to and directly following antigen exposure. Expression of *TBX21* (T-bet), an important transcription factor for atBC development, was also elevated in activated B cells in INS further supporting their bias for extrafollicular activation (**fig. 2E**).

Nevertheless, most of the INS-associated B cell states uncovered by differential abundance testing mapped to activated B cells [actNaïve (B3) and actMBC (B7)], with frequencies of these subsets being significantly elevated over both healthy and disease controls (**fig. 2D, F**). Control-associated states were overwhelmingly associated with naïve clusters, though many INS-associated states were also observed in these clusters denoting widespread dysregulation of the pre-antigenic B cell compartment (**fig. 2D**). To identify the pathways associated with dysregulation, we performed pseudobulk differential gene expression analysis between B cells from children with INS and healthy controls (**fig. S2D, data file S2**). The INS-associated B cell signature included genes associated with BCR signaling, antibody production, and type-I interferon signaling, with many differentially-expressed genes being prevalent across naïve, activated, efMBC, and foMBC populations (**fig. S2E, data file S3**) (*14*). All major genes involved in BCR signal transduction (*CD19*, *CD79B*, *SYK*, *BTK*, *BLNK*, and *PLCG2*) were particularly upregulated in actNaïve and actMBC subsets in INS, denoting a primed, or hyperactive, phenotype (**fig. 2G, H**). Activated subsets in INS also showed increased expression of genes involved in antigen presentation by MHC-II, denoting a greater capacity to receive T cell help and, accordingly, potentiate BCR signaling (**fig. 2H**). Accordingly, regulon assessment using the collecTRI resource revealed that NFκB activity, a major signaling endpoint downstream the BCR, was elevated in activated B cells from INS over healthy controls (**fig. 2I**) (*27*). Indeed, NFκB target genes *TNF* and *ICAM1* were amongst the most differentially expressed genes in INS (**fig. 2J**). *NR4A1*, encoding the transcription factor Nur77, the expression of which is directly associated with BCR signal strength, was highly expressed by activated B cells in INS altogether indicating that the hyperactive transcriptional profile translates to more potent BCR signaling (**fig. 2J**) (*28*). Since strong BCR signals favour extrafollicular responses over germinal center formation, we assessed whether the extrafollicular bias in INS was linked to enhanced BCR signaling. Evidence of increased BCR signaling was also observed in atBCs and MZ-like B cells, both of which expressed greater levels of BCR signaling genes, NFκB targets, and *NR4A1* (**fig. 2J, K**). IRF4 activity, which is driven by NFκB signaling and promotes plasmablast formation through the extrafollicular route, was strongly induced in these populations in INS over healthy controls (**fig. 2L**). Indeed, *BCL6* expression, a key driver of entry into germinal center responses that is repressed by IRF4, was diminished in activated B cells and MZ-like B cells in INS (**fig. 2M**). *In vitro* BCR stimulation of B cells validated these results, with B cells from children with INS exhibiting significantly greater rates of conversion into plasmablasts than those from healthy controls (**fig. 2N**). Collectively, these data demonstrate that B cells in INS are primed for strong BCR signaling, which in turn favours extrafollicular responses leading to atBC expansion and plasmablast formation.

### V_H_4-39^+^ clones are highly prevalent and uncensored from the plasmablast compartment in INS

B cell hyperactivity frequently reflects impaired central or peripheral B cell tolerance, in turn, permitting autoreactive clones to persist in the host. In this setting, clones that should be tolerized or deleted instead gain functional competency ultimately creating a skewed mature B cell repertoire enriched with autoreactive cells (*29, 30*). The B cell repertoire in INS remains poorly characterized. We predicted that B cell hyperactivity in INS would result in a biased repertoire. Assessment of BCR isotypes in INS showed decreased *IGHD* utilization across naïve (B0, B1, B9) and actNaïve (B3) B cell subsets in INS with compensatory use of *IGHM* (**fig. 3A, fig. S3A**). This decreased *IGHD* was similarly observed in the disease control and likely reflects the hyperactive B cell state (*31*).

**Figure 3.**
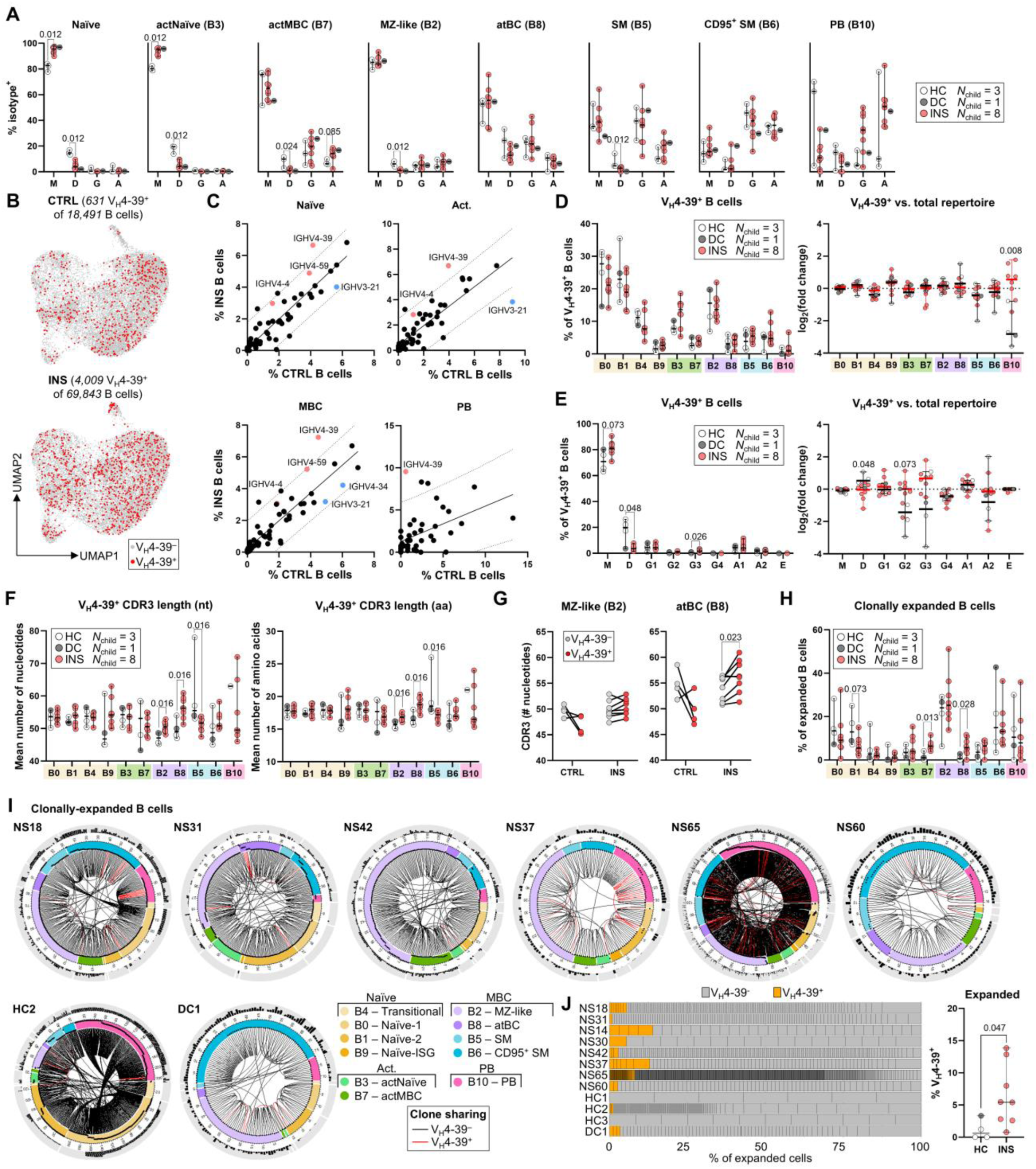
V_H_4-39^+^ clones are prevalent and persist into the plasmablast compartment in INS. (**A**) Isotype frequencies, based on *IGHC* usage, for B cell clusters from children with INS, HCs, and DC. (**B**) UMAP of the total B cells with V_H_4-39^+^ cells depicted in red. (**C**) Correlation of *IGHV* gene usage between INS and controls for each B cell metacluster. Significantly upregulated or downregulated genes are highlighted in orange and blue, respectively. A simple linear regression line with 95% prediction bands is provided for visualization. (**D**) B cell cluster frequencies in the V_H_4-39^+^ repertoire (*left*) and their log_2_(fold change) from cluster frequencies in the total repertoire (*right*). (**E**) Isotype frequencies within the V_H_4-39^+^ repertoire (*left*) and their log_2_(fold change) from isotype frequencies in the total repertoire (*right*). (**F**) Complementarity-determining region (CDR) 3 nucleotide (nt) and amino acid (aa) length in V_H_4-39^+^ cells from each cluster. (**G**) CDR3 nucleotide length in MZ-like B cells and atBCs compared between V_H_4-39^+^ and V_H_4-39^−^ cells. (**H**) B cell cluster frequencies within the clonally expanded pool. (**I**) Circos plots of clonally expanded B cells from the six children with INS and two control children showing high levels of clonal expansion. Cells are arranged by cluster from highest to lowest degree of clonal expansion (*clockwise*). The outer ring shows somatic hypermutation expressed as the percentage of mutated nucleotides in *IGHV* with the ring thickness representing 20% mutated nucleotides. Lines in the middle link clonal families with red lines denoting V_H_4-39^+^ clones. (**J**) Stacked bar plots with each stack representing a single clonal family (*left*) and frequencies of V_H_4-39^+^ cells within the expanded pool (*right*). In A, C, D, E, F, G and H, each dot represents a single donor with summary statistics represented as median with 95% confidence intervals. *P*-values were determined using independent Mann-Whitney *U*-tests comparing INS and healthy controls (A) or INS and controls (C, D, E, F, H), and independent Wilcoxon signed rank tests comparing V_H_4-39^−^ and V_H_4-39^+^ cells (G).

B cells in INS were significantly enriched for BCRs using *IGHV4-39* and *IGHV4-59* heavy chains, which were previously associated with autoreactivity in other systemic autoimmune conditions (**fig. 3B, C, fig. S3B**) (*29, 32, 33*). Of the two, V_H_4-39^+^ B cells (using *IGHV4-39*) were significantly enriched across naïve, activated, and memory compartments, and even persisted into the plasmablast pool, denoting a selection advantage for V_H_4-39^+^ clones in INS (**fig. 3C, fig. S3C**). While the overall frequency of VH4-39⁺ B cells did not differ between children with INS and controls, analysis of B-cell subset composition within the VH4-39⁺ repertoire revealed differences in selection (**fig. 3D**). When normalized to the complete B-cell repertoire (expressed as log₂ fold change), VH4-39⁺ clones were relatively enriched in the plasmablast compartment in children with INS, consistent with reduced censoring of these clones (**fig. 3D**) (*33*). V_H_4-39^+^ B cells in INS had diminished *IGHD* use favouring *IGHM*, whereas V_H_4-39^+^ clones in controls were better retained within the IgD^+^ compartment (**fig. 3E**). Notably, greater CDR3 lengths, a characteristic linked to antibody polyreactivity and autoreactivity, were observed in V_H_4-39^+^ atBCs and MZ-like B cells in INS than in healthy controls (**fig. 3F**) (*29, 34, 35*). V_H_4-39^+^ atBCs in INS even exhibited longer CDR3 lengths than V_H_4-39^−^ atBCs from the same individual, suggesting a possible relationship between V_H_4-39 BCRs and atBCs in INS (**fig. 3G**). Gene usage at *D*, *J*, and *L*/*K* loci were comparable between INS and controls (**fig. S3D-E**).

We observed no major differences in clonality between INS and controls, with six of the eight children with INS and two of the four controls showing high clonal expansion (defined as ≥5 B cells/clone) (**fig. S4A-D**). Nevertheless, clonally expanded B cells in INS contained more atBCs and actMBCs suggesting that extrafollicular B cell expansion is antigen directed (**fig. 3H**). Isotype use was not significantly different between clonally expanded B cells in INS and controls (**fig. S4E**). Importantly, we observed that V_H_4-39^+^ clones were present in the clonally expanded pool in INS and were largely absent from controls, again suggesting that V_H_4-39^+^ clones in INS are breaching tolerance mechanisms that are otherwise present in healthy individuals (**fig. 3I, J**). Together, these findings illustrate a skewed mature B cell repertoire in INS that may support the activation of clones carrying autoreactivity-prone BCRs.

### Autoreactivity against the slit diaphragm extends to CRB2

While autoantibodies to Nephrin are now well-documented in idiopathic podocytopathies, recent reports have shown that the underlying autoimmunity may extend to closely related slit diaphragm proteins like Kirrel1 and Podocin. We sought to characterize the seroprevalence of autoantibodies against CRB2, which has previously been reported in mouse, but not in humans, relative to anti-Nephrin in a cohort of individuals with active idiopathic podocytopathies (*N* = 82, including all patients analyzed above) (**fig. 1A, table S1**). People with other glomerulopathies (*N* = 22), and healthy individuals (*N* = 28) were used as controls. While anti-Nephrin IgG was prevalent in the cohort, anti-CRB2 IgG prevalence was markedly higher (**fig. 4A**): 20/42 (48%) children with SSNS/MCD, 3/9 (33%) children with SRNS/primary FSGS, 9/10 (90%) adults with MCD, and 18/21 (86%) adults with primary FSGS. Only small fractions of individuals with membranous nephropathy or other glomerulopathies carried either anti-Nephrin or anti-CRB2 denoting specificity to idiopathic podocytopathies (**fig. 4A**).

**Figure 4.**
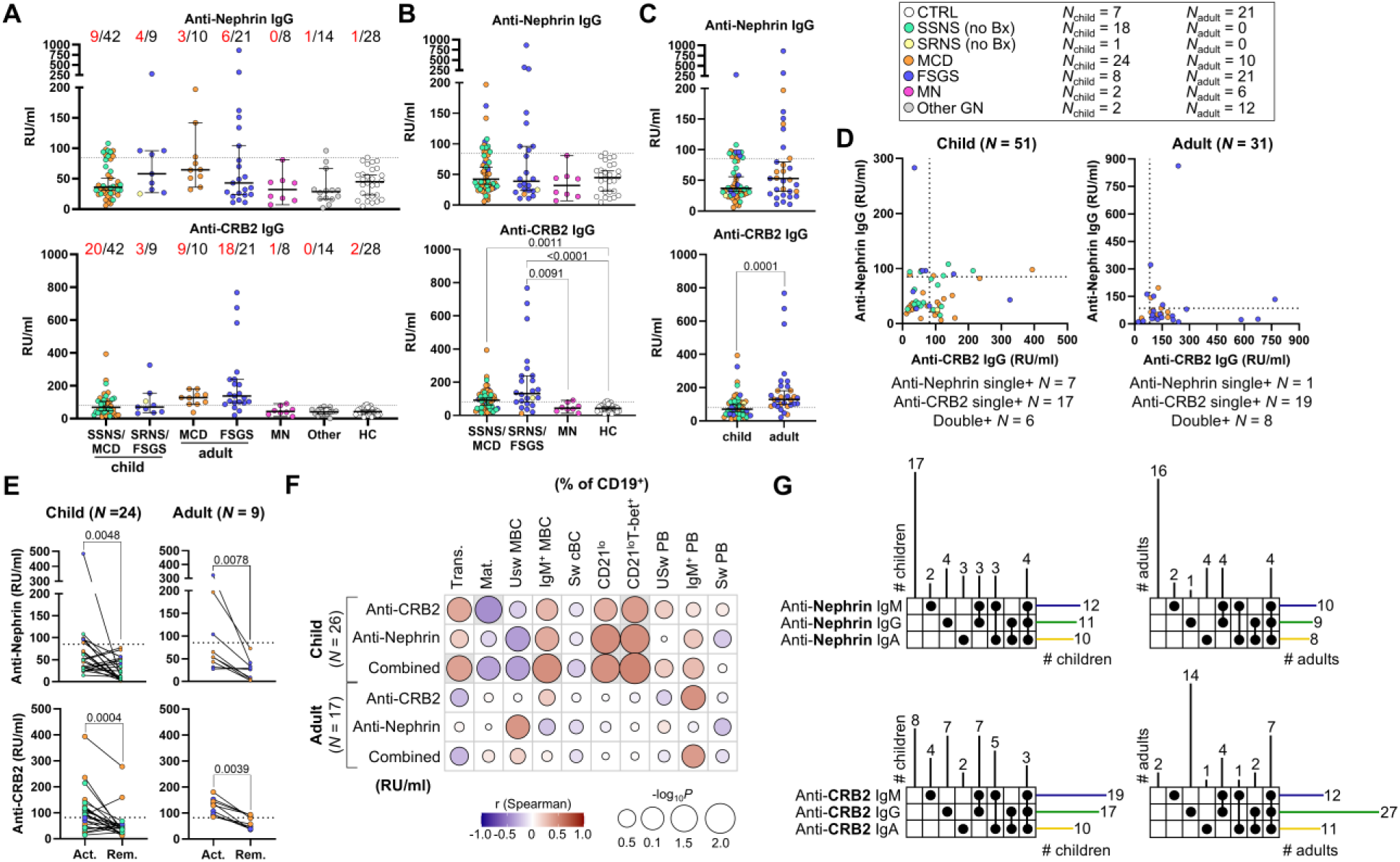
Anti-CRB2 autoantibodies are prevalent in idiopathic podocytopathies and can co-occur with anti-Nephrin autoantibodies. (**A**) Profiling of anti-Nephrin IgG and anti-CRB2 IgG in sera of patients with INS (*N*_child_ = 21), including SSNS (*N*_child_ = 20) and SRNS (*N*_child_ = 1), or MCD (*N*_child_ = 22, *N*_adult_ = 10), primary FSGS (*N*_child_ = 8, *N*_adult_ = 21), and membranous nephropathy (MN, *N*_child_ = 2, *N*_adult_ = 6), or other glomerular diseases (Other GN, *N*_child_ = 2, *N*_adult_ = 12). The seropositivity threshold determined using sera from HCs (*N*_child_ = 7, *N*_adult_ = 21) is depicted as a dotted line. (**B**) Anti-Nephrin IgG and anti-CRB2 IgG titers for idiopathic podocytopathies divided by steroid responsiveness and histological diagnoses, compared to MN and HC. **(C)** Anti-Nephrin IgG and anti-CRB2 IgG are compared between children and adults with idiopathic podocytopathies. **(D)** Correlations between anti-Nephrin IgG and anti-CRB2 IgG titers in children and adults with idiopathic podocytopathies. **(E)** Changes in anti-CRB2 IgG and anti-Nephrin IgG titers during active disease and remission in children (*N* = 24) and adults (*N* = 9) with idiopathic podocytopathies. **(F)** Correlation between frequencies of various B cell subsets and titers for anti-Nephrin IgG, anti-CRB2 IgG, and both autoantibodies combined in children (N=26) and adults (N=17) with idiopathic podocytopathies. **(G)** Upset plots depicting seropositivity patterns for anti-CRB2 and anti-Nephrin IgG, IgM and IgA in children (N=36) and adults (N=32) with idiopathic podocytopathies. In A-E, each dot represents a single patient and summary statistics are expressed as a median with 95% confidence intervals. *P*-values were determined using Kruskal-Wallis tests with Dunn’s correction for multiple comparisons in A and B, Mann-Whitney *U*-tests in C and E, and non-parametric Spearman correlations in F.

The highest anti-Nephrin and anti-CRB2 titers were observed in individuals with primary FSGS (**fig. 4B**). Notably, while anti-Nephrin IgG seropositivity was comparable in children and adults (9/31 [29%] adults, 13/51 [23%] children, *P* = 0.6136), anti-CRB2 IgG seropositivity was more strongly associated with adult idiopathic podocytopathies (27/31 [87%] adults, 23/51 [45%] children, *P* = 0.0002) suggesting that anti-CRB2 autoantibodies may arise from more sustained podocyte-directed B cell responses (**fig. 4A**). Accordingly, anti-CRB2 IgG titers were greatest in affected adults and seropositivity was significantly associated with onset during adulthood (**fig. 4C, fig. S5A**). Adults also showed greater incidence of co-seropositivity than children (8/31 [26%] adults, 6/51 [12%] children, *P* = 0.1330), suggesting longer-standing autoimmunity in adults. Nevertheless, these co-seropositivity measurements likely underestimate loss of tolerance to both antigens as longitudinal assessments show variable changes in seropositivity with time (**fig. S5B**).

Both autoantibodies correlated with disease activity, with decreased titers in remission likely associated with immunosuppressive treatments (**fig. 4E**). Between the two, anti-Nephrin IgG seropositivity was associated with higher proteinuria in both children and adults, while anti-CRB2 IgG did not correlate with proteinuria (**fig. S5C, D**). In summary, we demonstrate that autoreactivity at the slit diaphragm extends to CRB2 with loss of tolerance to both antigens being commonplace within the same individual.

### Autoantibodies correlate with extrafollicular responses in children with INS

We next sought to examine whether loss of tolerance to slit diaphragm proteins was associated with extrafollicular responses in children and adults with idiopathic podocytopathies. Anti-Nephrin and anti-CRB2 IgG titers correlated significantly with frequencies of CD21^low^ T-bet^+^ atBC in affected children, and the strength of this correlation improved when considering both autoantibodies together (**fig. 4F, fig. S6A**). This was also observed with IgM^+^ MBCs, which have previously been shown to be comprised mainly of recirculating MZ B cells (**fig. 4F**). Autoantibody titers in adults, however, showed minimal correlation with any B cell subset, and no clear correlations were observed with plasmablasts in either children or adults (**fig. 4F, S6A, B, S7A, B**).

Since presence of IgM antibodies is usually indicative of extrafollicular responses, we tested the isotype of slit diaphragm autoantibodies as confirmation. In children, seropositivity for IgM forms of anti-Nephrin and anti-CRB2 was comparable to IgG indicating that extrafollicular responses against these antigens are taking place at onset/relapse (**fig. 4G**). In adults, while anti-Nephrin IgM seropositivity was again comparable to IgG, anti-CRB2 IgG was the predominant form of anti-CRB2 antibody (**fig. 4G**). Altogether, this data provides evidence that podocyte-directed autoantibody responses emerge through extrafollicular responses.

### Epitope spreading at the slit diaphragm supports autoantibody development

We next assessed whether anti-CRB2 autoantibodies localized with podocytes in individuals with idiopathic podocytopathies, as was previously observed with anti-Nephrin. Anti-CRB2 IgG seropositive individuals (*N* = 5 with flash-frozen biopsy tissue available) showed punctate IgG deposits in glomerular podocytes, including in those who were concomitantly seronegative for anti-Nephrin IgG (**fig. 5A, fig. S8A, B**). In contrast, IgG staining in individuals seronegative for either autoantibody was negligible (*N* = 2). High resolution microscopy of a donor seropositive for both anti-CRB2 and anti-Nephrin IgG showed that IgG overlapped with both CRB2 and Nephrin, with certain zones showing co-localization with only one protein and not the other, consistent with the presence of anti-CRB2 autoantibody binding to podocytes and underlining their possible role in pathogenesis (**fig. 5B**).

**Figure 5.**
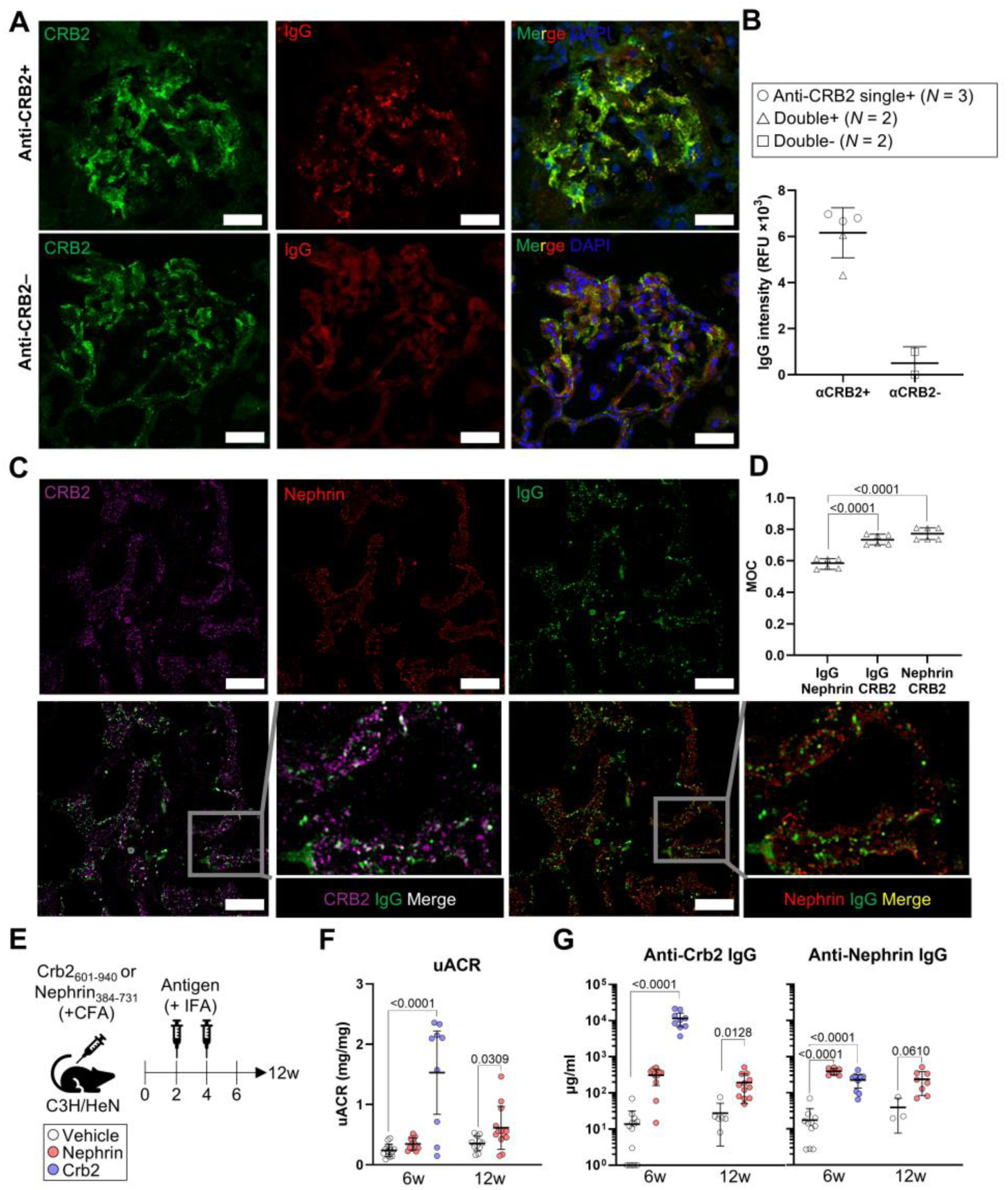
Epitope spreading at the slit diaphragm promotes autoantibody emergence. **(A)** Representative glomeruli from flash frozen kidney biopsies from anti-CRB2 IgG seropositive and seronegative patients with idiopathic podocytopathies showing CRB2 (*green*) and IgG (*red*) localization. **(B)** Average IgG staining intensity across all glomeruli are compared between anti-CRB2 IgG seropositive (*N* = 5) and seronegative (*N* = 2) individuals. **(C)** Representative glomerulus from a single patient with both anti-CRB2 and anti-Nephrin IgG showing CRB2 (*magenta*), Nephrin (*red*), and IgG (*green*) co-localization visualized with high-resolution structured illumination microscopy. **(D)** Overlap between IgG and Nephrin, IgG and CRB2, and Nephrin and CRB2 expressed as the Manders’ overlap coefficient (MOC) across three different areas of two glomeruli from the patient in H. **(E)** Schematic showing mouse immunization schedule for inducing experimental autoimmune nephrotic syndrome. **(F)** Urinary albumin-to-creatinine ratios (uACR) from immunized mice at 6-weeks and 12-weeks following immunization. **(G)** Anti-CRB2 IgG and anti-Nephrin IgG titers in immunized mice. In B, each dot represents a single patient, in D, each dot represents a distinct region of a glomerulus, and in F and G, each dot represents a single mouse. Summary statistics are expressed as a median with 95% confidence intervals. *P*-values were determined using one-way ANOVA with Tukey’s correction for multiple testing (D, F, G).

Nephrin and CRB2 are intimately proximal within the slit diaphragm. Consequently, we hypothesized that the break in tolerance to both autoantigens might arise by epitope spreading – the process by which the scope of an immune response gradually expands to recognize additional epitopes on the same or physically-associated antigens within the broadly dysregulated B cell milieu (*36*). To test this, we immunized mice with a portion of the extracellular domains of either mouse Crb2 (Crb2_601-940_) or Nephrin (Nephrin_384-731_) and re-challenged them at 14 and 28 days following initial immunization to induce podocytopathy (**fig. 5E**). Immunization with either autoantigen elicited proteinuria, though this was more profound with Crb2 immunization (**fig. 5F**). And as expected, both groups generated robust autoantibody responses to the immunizing antigen (**fig. 5G**). Strikingly, we also observed significant induction of autoantibodies against the non-immunizing antigen in both settings, indicating that epitope spreading within the slit diaphragm may contribute to the diversification of autoantibody species in idiopathic podocytopathies (**fig. 5G**).

## DISCUSSION

The recent discovery of autoantibodies targeting slit diaphragm proteins in INS, MCD, and FSGS supports a shared autoimmune basis for these idiopathic podocytopathies. However, the immune pathways that give rise to autoreactivity against podocyte antigens have remained largely undefined. In this study, we identify a consistent expansion of CD21^low^ T-bet^+^ atBCs across idiopathic podocytopathies, implicating this population in disease-associated humoral immunity. In children with INS, the expansion of atBCs is most consistent with extrafollicular B cell responses driven by strong B cell receptor signaling, whereas in adults, additional engagement of follicular responses was observed. We further demonstrate that slit diaphragm-directed autoreactivity extends to CRB2 and that autoantibodies against CRB2 and Nephrin frequently arise together, in particular in adults. Notably, the correlation between autoantibody titers and atBC abundance in pediatric INS supports an extrafollicular pathway as a plausible route by which podocyte-reactive B cells enter the antibody-secreting cell compartment.

CD21^low^ T-bet^+^ atBCs are frequently observed in conditions with chronic antigen exposure, including autoimmunity, infection, and immunodeficiency (*24, 25, 37*). In systemic lupus erythematosus, for example, atBCs are enriched with autoreactive clones and participate in anti-nuclear autoantibody production through extrafollicular conversion into plasmablasts (*17, 18*). Recently, we described CD21^low^ T-bet^+^ atBC expansion in children with INS, which has since been confirmed by other studies (*14, 38, 39*). Here, we extended this finding to adults with MCD/FSGS. While CD21^low^ T-bet^+^ atBC expansion is insufficient to assert the presence of extrafollicular responses (*40*), transcriptional work demonstrated that B cells in INS were preferentially enriched in states associated with extrafollicular responses. These were defined by low somatic hypermutation, class-switch recombination, and expression of key extrafollicular response genes (e.g., *GPR183*, *TNFRSF13B*, and *TBX21*) (*19, 26*). In children with INS, we also observed that frequencies of circulating atBCs correlated with anti-Nephrin and anti-CRB2 titers positioning the extrafollicular response as a relevant route into the antibody-secreting compartment for podocyte-reactive clones. Supporting an extrafollicular origin, anti-Nephrin and anti-CRB2 IgM autoantibodies were more prevalent than either IgG or IgA in children with INS. The expansion of atBCs and the induction of extrafollicular responses may reflect chronic antigen exposure, which is exemplified by a relapsing-remitting disease course.

We did not find any correlation between atBCs and autoantibodies in adults with MCD/FSGS. B cell profiling by flow cytometry suggested more profound B cell dysregulation in adults than children. Mature naïve B cells were significantly reduced whereas the pool of isotype-switched CD27^+^ CD21^+^ MBCs was enlarged. This MBC population largely comprises germinal center-derived B cells and may hence reflect the entry of podocyte-reactivity into the germinal center-dependent arm of B cell responses (*23*). Podocyte-reactivity in the long-lived plasma cell compartment, whose emergence is dependent on germinal centers, may explain the greater propensity for steroid resistance and diminished effectiveness of rituximab in adults with FSGS (*20, 21*). This germinal center-dependent origin for autoantibodies would also explain the lack of correlation with circulating B cell subsets in adults. In contrast, podocyte-reactivity in the short-lived plasmablast pool arising from extrafollicular responses in childhood INS may explain the greater likelihood of steroid-sensitivity and increased success of CD20-based B cell depletion (*21*). Indeed, children with INS showed elevated frequencies of plasmablasts whereas this was only observed in some adults. Nevertheless, direct demonstration of podocyte-reactivity in circulating B cells remains lacking in idiopathic podocytopathies and is a focus of our future endeavours.

We sought to better understand the mechanisms driving the extrafollicular skewing of B cells in childhood INS. Strong BCR signals, in combination with interferon and TLR signaling, promote extrafollicular B cell activation while moderate BCR signals promote germinal center formation (*41*). Single-cell RNA-sequencing revealed that the B cell compartment in INS was enriched in pre- and post-antigenic B cells displaying an activated phenotype (actNaïve and actMBC). B cells with similar transcriptional profiles have been previously described in humans with high variability between individuals and therefore likely reflect a cell state rather than a true B cell subset (*42*). These cells expressed high levels of genes involved in BCR signaling including *CD19*, *CD79B*, *SYK*, *BTK*, *BLNK1*, and *PLCG2*. This hyperactive phenotype likely augmented BCR signaling as was denoted by increased *NR4A1* expression in INS B cells. Accordingly, greater NFκB activity was observed in INS B cells alongside IRF4, a major determinant of plasmablast differentiation. *In vitro* testing confirmed a greater propensity for plasmablast differentiation by INS B cells following BCR stimulation. This hyperactive B cell phenotype is likely a consequence of underlying genetics (*10*) as several predisposing polymorphisms may either directly act to enhance BCR signaling (e.g. *PLCG2*, the expression of which was highly elevated in INS B cells) or indirectly by promoting T-B interactions (e.g., HLA-II polymorphisms, *CLEC16A*, and *CD28*) (*43–47*).

In the context of B cell hyperactivity, reduced activation thresholds may enable autoreactive B cell clones to evade tolerance checkpoints. In autoimmunity, this leads to a skewed BCR repertoire enriched for autoreactive specificities, with preferential usage of certain V_H_4 gene segments, including V_H_4-34, V_H_4-39, and V_H_4-59, which are commonly expanded in autoimmune settings (*29*). In children with INS, we observed an important expansion of V_H_4-39^+^ clones. While these clones seemed to be better excluded from both clonally expanded B cell and plasmablast pools in controls, V_H_4-39^+^ clones were readily present in both compartments in INS. This parallels a recent study in anti-phospholipid syndrome showing that V_H_4-39^+^ antiphospholipid-specific clones persist from the naïve repertoire into the MBC and antibody-secreting cell repertoires to a greater extent than in healthy individuals (*33*). Furthermore, V_H_4-39 BCRs have also been linked with polyreactivity (*48*). Indeed, V_H_4-39^+^ atBCs in INS had longer CDR3 sequences, a feature associated with polyreactivity, than V_H_4-39^+^ atBCs in controls and V_H_4-39^−^ atBCs from the same individual (*29, 34, 35*). While these findings support the idea that B cell hyperactivity promotes the expansion of potentially autoreactive and polyreactive clones in INS, the specificity of these clones has not been tested. Nevertheless, two previous studies in children with INS and adults with membranous nephropathy reported increased *IGHV4-39* utilization by B cells, hence supporting a possible link between V_H_4-39 BCRs and podocyte autoreactivity (*49, 50*).

Autoantibodies against the slit diaphragm proteins Nephrin, Podocin, and Kirrel1 have been described in INS (*5, 6, 51*). Here, we show that autoreactivity at the slit diaphragm extends to CRB2, a critical protein for glomerular filtration that regulates Nephrin positioning (*52, 53*). Anti-CRB2 was prevalent in both children and adults with idiopathic podocytopathies but showed greater accumulation in adults, in contrast with anti-Nephrin, which showed similar seroprevalence in children and adults. This suggests that anti-CRB2 may be increasingly associated with long-standing podocyte-reactivity. We observed a greater degree of non-IgM isotypes for anti-CRB2 autoantibodies in adults, in agreement with our hypothesis that podocyte-reactivity in adults may have progressed into the germinal center-dependent arm that promotes class-switch recombination. Consistent with previous studies, we observed that anti-Nephrin seropositivity was associated with proteinuria (*6, 54*) unlike anti-CRB2, though titers of both autoantibodies decreased substantially in remission. While we provide evidence that anti-CRB2 antibodies bind to podocytes in affected individuals, the relative contribution of each autoantibody to podocyte injury remains to be determined. Association of anti-Nephrin, however, with post-transplant recurrence of MCD/FSGS does support a pathogenic role for the autoantibody itself (*5, 55, 56*).

Importantly, we observed significant overlap in anti-Nephrin and anti-CRB2 seropositivity in both children and adults with idiopathic podocytopathies. The marked temporal variability in anti-CRB2 and anti-Nephrin seropositivity within individuals, potentially driven by immunosuppressive therapy or disease progression, suggests that our cohort underestimates the prevalence of Nephrin- and CRB2-directed autoimmunity. This is consistent with a recent study showing that up to 90% of children with INS carry anti-Nephrin autoantibodies during active disease in the absence of previous immunosuppression, with much lower seroprevalence observed in relapses following exposure to immunosuppressive treatment (*6*).

Using a mouse model, we demonstrated that loss of tolerance to both Nephrin and CRB2 can result in intermolecular epitope spreading, providing an explanation for the co-occurrence of autoantibodies against both antigens. This process, which often takes years to progress, is a well-established feature of autoimmune diseases and can correlate with clinical severity, as is the case in systemic lupus erythematosus where epitope spreading drives the diversification of autoantibodies targeting ribonucleoprotein complexes (*36, 57*). Notably, MCD and FSGS can also occur secondary to systemic lupus erythematosus (*58, 59*). It has recently been shown that CD21^low^ T-bet^+^ atBCs can mediate epitope spreading through antigen presentation to CD4^+^ T cells (*60*). Epitope spreading between slit diaphragm proteins would suggest B cell access to protein complexes, for example through tissue damage or the release of extracellular vesicles (*61, 62*). While our findings are compatible with epitope spreading as a downstream consequence of an initial breach of tolerance to slit diaphragm antigens, definitive evidence for this temporal sequence is still lacking. To this end, longitudinal studies tracking the evolution of the podocyte-directed autoantibody response from disease onset are needed.

Beyond epitope spreading, the high prevalence and IgG-dominant profile of anti-CRB2 antibodies, particularly in adults, suggest that CRB2 autoreactivity may reflect more sustained or mature podocyte-directed immunity. Their frequent presence in individuals lacking anti-Nephrin antibodies indicates that CRB2 is an independent target of humoral autoreactivity, raising the possibility that anti-CRB2 could serve as a complementary biomarker for chronic or diversified immune activity. Given that adult disease often shows reduced responsiveness to CD20-based therapies, the predominance of anti-CRB2 IgG may also point to a greater contribution of long-lived or germinal center-derived plasma cells, with potential implications for therapeutic stratification.

This study has some limitations. Owing to the rarity of idiopathic podocytopathies, cohort sizes were necessarily modest, and paired PBMC samples were not available for all individuals with corresponding serum samples. Although our cohort is ethnically diverse and broadly representative of individuals with idiopathic podocytopathies in Canada, validation in independent international cohorts will be important. In addition, most samples were obtained during relapse rather than at disease onset, and many individuals had received immunosuppressive therapy, which may have attenuated autoantibody seropositivity and plasmablast frequencies. Transcriptomic analyses were restricted to children with steroid-sensitive nephrotic syndrome or biopsy-proven minimal change disease in order to capture early disease features while limiting the confounding effects of immunosuppression; nevertheless, extension of these analyses to other podocytopathies and disease stages will be informative. Finally, we did not directly assess podocyte antigen-specificity within B cell populations, which could be addressed in future studies through antibody cloning or antigen tetramer-based approaches.

In summary, we identify extrafollicular, hyperactivation-driven B cell responses as a plausible pathway by which podocyte-reactive clones escape immune tolerance. Importantly, we expand the landscape of slit diaphragm-directed autoimmunity to include CRB2 and support intermolecular epitope spreading as a potential mechanism underlying the diversification of podocyte-targeting autoantibodies. Together, these findings provide a mechanistic immune framework that may inform the rational development of B cell-targeted therapies and immune-based biomarkers for idiopathic podocytopathies.

## MATERIALS AND METHODS

### Study design

Blood from 104 patients with glomerulopathies and 48 healthy controls were collected and used in this study (**table S1**). Of these, 75 patients and 44 healthy controls were enrolled in the McGill University Health Centre Kidney Disease Biorepository (MUHC-14-466, T.T.), while 29 patients and 4 healthy controls were enrolled in the CHILDNEPH study biobank (CHREB-16-2186, S.S.) (*14, 63*). In total, there were 55 children enrolled in the study, 19 of whom were non-biopsied and classified as having SSNS (*N*=18) or SRNS (*N*=1) while the remaining were diagnosed through biopsy with MCD (*N*=24), primary FSGS (*N*=8), membranous nephropathy (*N*=2), or other glomerulopathies (*N*=2) (**table S1**). All 49 adults had biopsy-proven disease, including MCD (*N*=10), FSGS (*N*=21), membranous nephropathy (*N*=6), and other glomerulopathies (*N*=12) (**table S1**).

Children were considered in active disease when proteinuria at time of blood draw exceeded 0.25 g/mmol on the urinary dipstick or 1 g/day with other measurements, and a diagnosis of either first episode or relapse was made by a nephrologist in a contemporary clinical evaluation. Adults were considered in active disease when proteinuria at time of blood draw exceeded 1 g/day and a diagnosis of either first episode or relapse was made by a nephrologist in a contemporary clinical evaluation. For children, remission was defined by a negative or trace urinary dipstick measurement or proteinuria less than 200 mg/day, while for adults it was defined by a proteinuria of less than 300 mg/day (*64*). For children, steroid resistance was defined by a persistent proteinuria of >200 mg/day despite four weeks of daily steroid therapy (equivalent of prednisone at 60 mg/m^2^/day) and a two week confirmation period with escalated steroid treatment, while in adults it was defined by a persistent proteinuria of >3500 mg/day with a reduction of <50% from baseline with 16 weeks of steroid therapy (equivalent of prednisone at 1 mg/kg/day) (*64, 65*). Healthy controls were defined as patients without a diagnosis of kidney, autoimmune, allergic, hematologic or neoplastic disease. Serum samples were obtained from all participants and 28 of the healthy controls, while PBMC were available for 43 patients and 35 healthy controls (**table S1**).

Clinical data was collected retrospectively when available for each patient followed at the McGill University Health Centre using their electronic medical file. For patients enrolled in the CHILDNEPH cohort, limited clinical data was available through the CHILDNEPH registry. All participants and/or their legal guardians gave written informed consent to be enrolled in the study, and biospecimens were used in accordance with our standard operating protocol (MUHC-15-341, T.T.). Research ethics board approvals were obtained for all centres involved in this study.

### Mice

Wildtype C3H/HeN (Charles River Laboratories) were bred at the Research Institute of the McGill University Health Centre. All mice were raised in specific pathogen-free conditions, and male 8-week-old mice were used in this study. Experiments were performed according to the Canadian Council on Animal Care and to institutional guidelines at McGill University, and ethical guidelines approved by local committees at the RI-MUHC.

### B cell profiling by flow cytometry

Cryopreserved PBMC were thawed in warm RPMI 1640 media (Wisent) completed with 10% FBS (Wisent), 10,000 U/ml penicillin-streptomycin (Gibco), 1% MEM nonessential amino acids (Wisent), 1% 1M HEPES solution (Wisent), and 1% 100 mM sodium pyruvate. Cells were rested at 37°C for 2 h in a 5% CO_2_ incubator before staining. A viability of over 90% was confirmed by Trypan blue, and a maximum of 1×10^6^ cells were transferred into a 96-well v-bottom plate for staining per flow cytometry panel. Cells were washed once in ice-cold PBS supplemented with 2% FBS and labelled with the Fixable Viability eFluor 780 dye (1:1000, ThermoFisher Scientific). After a 15-minute incubation at 4°C, cells were washed again in PBS+2% FBS, and incubated with Fc receptor block (1:50, BD Biosciences) for 15 minutes. Cells were washed and staining for extracellular markers was performed using antibody cocktails prepared in ice-cold PBS+2% FBS and Brilliant Stain Buffer (1:2, BD Biosciences): anti-human CD3ε BV785 (1:50, OKT3, BioLegend), anti-human CD19 BV605 (1:20, SJ25C1, BD Biosciences), anti-human CD20 AlexaFluor 700 (1:50, 2H7, BioLegend), anti-human CD21 BV421 (1:20, B-ly4, BD Biosciences), anti-human CD27 PE-Cy7 (1:20, M-T271, BD Biosciences), anti-human CD10 BUV737 (1:20 HI10a, BD Biosciences), anti-human CD38 BUV737 (1:20, HB7, BD Biosciences), anti-human CD11c PerCp-Cy5.5 (1:20, B-ly6, BD Biosciences), anti-human IgD BV510 (1:20, IA6-2, BioLegend), anti-human IgM AlexaFluor 488 (1:40, MHM-488, BioLegend), anti-human FcRL5 APC (1:20, 509f6, BioLegend), anti-human CD24 PE (1:20, ML5, BD Biosciences), anti-human CD1c BV711 (1:20, L161, BioLegend), anti-human CXCR5 APC (1:20, J252D4, BioLegend), anti-human TACI PE (1:20, 1A1, BioLegend), anti-human GPR183 PerCp-Cy5.5 (1:50, SA313E4, BioLegend), and anti-human CD23 APC (1:20, EBVCS-5, BioLegend). Cells were incubated at 4°C for 20 minutes, washed, and then fixed/permeabilized using the eBioscience Foxp3/Transcription Factor Staining Buffer Set (eBioscience). Cells were then washed with 1X permeabilization buffer (eBioscience) and stained for intracellular markers using antibody cocktails prepared in the same buffer: anti-Ki-67 BUV395 (1:50, B56, BD Biosciences), and anti-T-bet PE (1:20, 4B10, BioLegend). Two final washes were performed in permeabilization buffer and PBS+2% FBS before cells were acquired on the BD LSR Fortessa X-20. Data were analyzed using FlowJo v10.8 software (FlowJo, LLC).

### Single-cell RNA-sequencing

#### Library preparation and sequencing

Cryopreserved PBMC were thawed as described in the previous section. Cells were washed with ice-cold PBS+2% FBS and labelled with an antibody cocktail containing Fixable Viability eFluor 780 dye (ThermoFisher Scientific, 1:1000), anti-human CD19 BV605 (1:20, SJ25C1, BD Biosciences), anti-human CD3ε BV785 (1:50, OKT3, BioLegend), AlexaFluor 700 anti-human CD20 (1:50, 2H7, BioLegend) and anti-human CD10 BUV737 (1:20, HI10, BD Biosciences) prepared in PBS+2% FBS at 4°C for 15 minutes. After washing, cells were filtered through 70 μm mesh and total live mature B cells (Viability eFluor 780^−^ CD19^+^ CD10^low^ CD3ε^−^) were isolated via fluorescence-activated cell sorting using the BD FACSAria Fusion with >99% purity. B cells were washed in PBS+0.04% bovine serum albumin, concentrated to 1,500 cells/μL, and processed using the Chromium Next GEM Single Cell 5’ v2 (PN-1000263) and Chromium Single Cell Human BCR Amplification (PN-1000253) kits in three batches (*N*=4 samples/batch) targeting 10,000 to 15,000 captured cells. Complementary DNA was synthesized and Chromium Single Cell 5’ gene expression and V(D)J libraries were prepared according to manufacturer instructions. Libraries were sequenced on a NovaSeq6000 (Illumina) to a median sequencing depth of 30,000 reads/cell for gene expression libraries and 5,000 reads/cell for V(D)J libraries.

#### Preprocessing of gene expression data

FASTQ reads were aligned to the GRCh38 reference genome v1.2.0 (refdata-cellranger-GRCh38-1.2.0) and count matrices were generated using CellRanger v7.1.0. All preprocessing and quality control steps were performed using the Seurat v5.0.0 R package (*66*). Briefly, contaminating non-B cells expressing *CD3G*, *CD14*, *LILRA4* or hemoglobin genes were removed, and cells with <200 of >2700 features, >8% transcripts belonging to mitochondrial genes, or <14% transcripts corresponding to ribosomal protein genes were filtered out as doublets or nonviable cells. Mitochondrial genes, ribosomal genes, *MALAT1* and *XIST* were regressed from the dataset to prevent their impact on downstream clustering.

#### Dimensionality reduction and clustering

The reciprocal principal component analysis (PCA) method in Seurat v5.0.0 was used to generate B cell clusters that were uniformly present across individuals and batches. Expression data were normalized, and variable features were identified for each independent sample. Features that were consistently variable across samples were then selected (SelectIntegrationFeatures), scaled, and used to perform an initial PCA. The resulting principal components (PCs) were used to identify integration anchors (FindIntegrationAnchors) and integrate the data (IntegrateData). The integrated data was scaled and PCA was performed again. A uniform manifold approximation and projection (UMAP) was generated on *k*NN values calculated with 10 PCs, and Louvain clusters were generated at a resolution of 0.5. A single cluster of cells corresponding with contaminating natural killer cell were removed and a new UMAP was generated using 9 PCs and a resolution of 0.5.

#### Cluster annotations

Marker sets for each cluster were generated by running differential gene expression analysis between the query cluster and all other B cells using the FindAllMarkers function in Seurat v5.0.0 (**fig. S2C**). The resulting gene lists were filtered to only include genes expressed in at least 10% of cells and were subsequently used to assign cluster identities (**fig. 2A**). Broadly, naïve B cells were identified based on the expression of *BACH2*, *TCL1A*, *FCER2*, and *IL4R*; activated B cells (Act) using *CD69*, *FOS*, and *FOSB*; memory B cells (MBC) using *BCL2A1*, *IGHA1*, and *CD27*; and plasmablasts (PB) using *PRDM1*, and *JCHAIN*.

#### Trajectory inference

Activated and MBC clusters from the entire dataset were isolated and a new UMAP was learned on 10 PCs. Trajectories were inferred using Monocle3 v1.3.5 selecting the actNaïve cluster as the starting point (*67*). Normalized density plots of B cells from children with INS and controls were generated using the same UMAP and overlayed with a simplified version of the trajectories for visualization purposes.

#### Differential abundance testing

Overlapping cellular neighbourhoods reflective of distinct cell states were generated on the integrated UMAP embeddings using MiloR v1.10.0 (*68*). Here, a *k*NN graph was built using the buildGraph and makeNhoods function on randomly selected cells in the data set (5% of total), 8 PCs, and *k* = 50. These parameters were fine-tuned through manual inspection of neighbourhood sizes as was recommended through the standard MiloR workflow. Shifts in proportions of distinct neighbourhoods between children with INS and controls was tested using the testNhoods function with a spatial *FDR* adjusting for the overlap between neighbourhoods of < 0.1 considered as significant.

#### Differential gene expression, gene set enrichment analysis, and transcription factor regulon enrichment

To identify a robust set of genes that are differentially expressed between children with INS (*N* = 8) and healthy controls (*N* = 3), pseudobulk differential gene expression analysis was carried out using Muscat v1.13.1 (*69*). Pseudobulk data were generated by aggregating normalized and log-transformed gene counts at the sample and cell cluster levels using the aggregateData function. Differential gene expression was then performed using edgeR and differentially expressed genes were considered as those with a |log_2_(fold change)| < 0.65, *P*_adj_ < 0.05, and were expressed in at least 10% of cells. Gene set enrichment analysis (GSEA) was performed on unfiltered gene lists ordered by -log_10_(*P*-value) × sign[log_2_(fold change)] using the GSEA App (Broad Institute) referencing the Gene Ontology and Reactome databases for gene sets that contained at least 5 and no more than 500 genes. Scores for transcription factor regulons, obtained from the CollecTRI gene regulatory network, were inferred for every cell using decoupleR v2.9.7 (*27*). Scaled scores were used for comparisons of regulons between children with INS and healthy controls.

#### V(D)J sequence processing and gene annotation

Demultiplexed pair-end FASTQ reads were processed via CellRanger v7.1.0 and aligned to the GRCh38 human reference genome (refdata-cellranger-vdj-GRCh38-alts-ensembl-5.0.0). The resulting outputs were then processed using the Immcantation pipeline. The initial germline annotations were performed using IgBLAST v1.22.0 with the IMGT/V-Quest reference, and the outputs were converted to the adaptive immune receptor repertoire (AIRR) format using MakeDb.py from Change-O v1.3.0 (*70–72*). Only sequences with productive rearrangement and a CDR3 nucleotide sequence length that is a multiple of three were retained, and cells with no or multiple heavy chains were removed. If a cell had multiple light chains, the most abundant transcript was kept (or the first transcript listed in case of a tie). Finally, only cells present in the gene expression dataset following processing, quality control, and clustering steps were retained in the V(D)J data.

#### Clonal lineage inference

Clonal lineages were inferred for each donor on productive heavy chain sequences with hierarchical clustering using SCOPer v1.3.0 (*73–75*). Heavy chains with equivalent V and J genotypes and CDR3 lengths were parsed into partitions, and CDR3 nucleotide sequences within a 0.15 normalized Hamming distance from each other within each partition were defined as clones. Full-length consensus germline sequences were reconstructed for each clone from the IMGT/V-Quest reference using Dowser v2.1.0 (*76*). Clone networks were then generated using the Dandelion v0.3.5 python package on the inferred clones and visualized through Scanpy v1.9.8 (*77, 78*).

#### Somatic hypermutation and class-switch recombination

Somatic hypermutation was calculated as the frequency of nucleotide mismatches from the germline sequence of the CDR1, CDR2, FWR1, FWR2, and FWR3 segments of the variable heavy chain using SHazaM v1.2.0 (*72*). CSR was calculated as the frequency of cells with a constant heavy chain genotype that was not *IGHM* nor *IGHD*. The constant heavy chain genotype was obtained from the ‘c_call’ column of the CellRanger output.

### In vitro plasmablast differentiation

Cryopreserved PBMC were thawed as described in the flow cytometry section. Cells were washed with ice-cold PBS+2% FBS and labelled with an antibody cocktail containing the Fixable Viability eFluor 780 dye (ThermoFisher Scientific, 1:1000), anti-human CD19 BV605 (1:20, SJ25C1, BD Biosciences), anti-human CD3ε BV785 (1:50, OKT3, BioLegend), and AlexaFluor 700 anti-human CD20 (1:50, 2H7, BioLegend). After a subsequent wash, cells were resuspended in ice-cold PBS+2% FBS, filtered through 70 μm mesh, and live B cells (Viability eFluor 780^−^CD19^+^ CD20^+^ CD3ε^−^) were isolated via fluorescence-activated cell sorting using the BD FACSAria Fusion with >99% purity. Cells were washed with and brought up in complete RPMI 1640 medium. Duplicate 40,000 cells were plated in 96-well round-bottom plates and stimulated with recombinant human CD40L-Ig (1 μg/mL, Invivogen), F(ab’)2 goat anti-human IgM/IgG (100 ng/mL, ThermoFisher), IL-21 (50 ng/mL, BioLegend), and IL-4 (10 ng/mL, BioLegend). After 4-day culture, cells were harvested and stained for flow cytometric evaluation of plasmablast (CD27^+^ CD38^high^) differentiation.

### Human anti-CRB2 and anti-Nephrin ELISAs

Wells of ELISA plates (Nunc-Immuno Plate, MaxiSorp, 96-well, ThermoFisher) were coated overnight at 4°C with 1.00 µg of CRB2_597-943_ or 0.13 µg of Nephrin in 0.2 M sodium carbonate-bicarbonate solution. Antigens were the extracellular domain of human CRB2 (E597-V943) produced in *E. coli* or full-length human Nephrin (SinoBiological) produced in HEK293 cells (*79*). After overnight coating, plates were washed three times with washing solution (PBS+0.2% Tween) before blocking with blocking solution (PBS+2% BSA and 0.2% Tween) for 1 hour. Then, plates were loaded with 100 µL of standards and serum samples in duplicates. The standard curve was generated using seven serial dilutions of polyclonal rabbit anti-human CRB2 antibody (custom produced by MediMabs, Montreal, using human CRB2_597-943_ recombinant protein) or monoclonal sheep anti-human Nephrin antibody (RnD systems). Sera was diluted 1:1000 (anti-CRB2) or 1:500 (anti-Nephrin) in blocking buffer. After 2 hours of incubation, plates were washed three times and incubated for 1 hour with anti-rabbit IgG-HRP (anti-CRB2) or anti-goat IgG-HRP (anti-Nephrin) antibodies (Abcam) for the standard curve, and anti-human IgG-HRP antibodies (Abcam) for sera. Plates were then incubated with 100 µL of 3,3’,5,5’-tetramethylbenzidine substrate solution (Bethyl laboratories) for 1 minute (anti-CRB2) or 5 minutes (anti-Nephrin) before the reaction was stopped with 2 M H_2_SO_4_ (Sigma-Aldrich). The results were read with a TECAN plate reader at an absorbance of 450 nm and titers were calculated using a four-parameter logistic curve. Uncoated plates loaded with the same serum dilutions were run to detect background. As the background was <1% of detected optical density, it was ignored. Positivity cut-offs were determined as *Cutoff* = *x̄* + *SD* × *f*, where *x̄* is the mean titer the healthy control population (presumed seronegative) and *SD* is the standard deviation. The multiplier *f* is calculated as *f* = *t*_1-0.05_ + (1/*N*), where *t*_1-0.05_ is the critical *t*-value for an *α*-value of 0.05 and *N* is the size of the cohort.

### Confocal imaging of human flash frozen kidneys

Flash frozen kidney biopsies from seven patients were prepared by the McGill University Health Centre kidney pathology department. Kidneys were sectioned at 4 μm, air dried for 30 minutes, and subjected to treatment with 95% ethanol for 10 minutes at -20°C. Sections were then washed three times in PBS before fixing with 4% paraformaldehyde (ThermoFisher) for 15 minutes. After another three washes in PBS, blocking was performed with PBS containing 3% bovine serum albumin (Sigma-Aldrich) and 2% FBS (Wisent) for 1 hour. Staining with rabbit anti-CRB2 (1:100, Sigma-Aldrich), guinea pig anti-Nephrin (1:100, Progen), and mouse anti-human IgG (1:750, Abcam) was carried overnight at 4°C. Sections were washed again in PBS, and stained with anti-rabbit AlexaFluor 488 (1:1000, Cell Signaling) and anti-mouse AlexaFluor 555 (1:1000, Cell Signaling) for 1 hour at room temperature. Staining with DAPI (1:1000, ThermoFisher) was carried out afterwards for 15 minutes, and the sections were mounted with Aquamount (Fisher Scientific). Immunofluorescence images were captured with a Zeiss LSM780 confocal microscope using a 20X objective. High-resolution structured illumination microscopy images were captured using a Zeiss LSM880 SIM confocal microscope. Laser intensity was kept the same for all images across a single experiment. Images were processed and quantified using Zeiss Microscopy Software v3.12 (Carl Zeiss AG).

### Mouse model for autoimmune podocytopathy

Recombinant mouse Crb2_601-940_ or Nephrin_384-731_ produced in *E. coli* in PBS (1 mg/mL) was emulsified 1:1 with Complete Freund’s Adjuvant (Millipore Sigma), and 100 µL was injected bilaterally at the base of tail for a total dose of 100 µg (*80*). Mice were re-challenged after 2 and 4 weeks with 100 µg of antigen emulsified in Incomplete Freund’s Adjuvant (Millipore Sigma) injected subcutaneously at both flanks. Urine was collected manually at the indicated time points, and urinary albumin (Bethyl Laboratories) and creatinine (Cayman Chemical) concentrations were measured by ELISAs according to manufacturer instructions. The former was normalized to the latter to obtain urinary albumin-to-creatinine ratios. Serum was collected from saphenous veins.

### Mouse anti-Crb2 and anti-Nephrin ELISAs

ELISAs for anti-Crb2 and anti-Nephrin in mice were conducted as described for human ELISAs. Coating antigens were replaced for mCrb2_601-940_ or mNephrin_384-731_. Serum samples were diluted 1:1000, 1:5000, or 1:25000 to ensure that optical densities fall within the linear range of the ELISAs. Detection was carried out using anti-mouse IgG-HRP antibodies diluted 1:1000 in blocking buffer (Abcam).

### Statistical analysis

Statistical analyses were performed on R v4.4.0 or GraphPad Prism v10.2.2 software. Continuous variables are presented as median with 95% confidence intervals with each point representing a single individual, while categorical variables are presented as number of patients (percentage). *P*-values for differences in categorical variables were determined using a Fisher’s exact test. Two-sided analyses between two unpaired groups were completed using Mann-Whitney U-tests, while paired groups were analyzed using Wilcoxon signed-rank tests. Comparisons between more than two groups were performed using Kruskal-Wallis tests and corrected with Dunn’s post-hoc testing, or on-way ANOVA corrected with Tukey’s post-hoc testing. Non-parametric Spearman correlations were used to obtain *r* and *P*-values when evaluating the relatedness of two parameters. For differential gene expression, gene set enrichment analysis, and pathway analysis, *P*_adj_-values were determined using Wilcoxon rank-sum tests with Benjamini-Hochberg corrections. *P*-values < 0.05 were considered significant. No sample size calculations were conducted *a priori*, and no samples were excluded from the study. Serum samples were available for some individuals without associated PBMC and *vice versa*.

## DATA AVAILABILITY

Raw sequencing data and processed count matrices were deposited in the NCBI Gene Expression Omnibus (GEO) under the accession number GSE307080. The results of all differential gene expression analyses are included in the Supplementary Data files of this paper. Public data repositories used include the Gene Ontology and Reactome databases accessed by the GSEA app and CollecTRI for transcription factor regulon assessment. The GRCh38 human reference genome was used for gene expression sequence alignment. The IMGT/V-Quest human reference was used for V(D)J sequence alignment.

## ACKNOWLEDGEMENTS

We would like to thank all study participants, their families, researchers, physicians, and personnel involved in the McGill University Health Centre Kidney Disease Biorepository and the CHILDNEPH biobanks. We also thank I. Ragoussis, H. Djambazian, A. Harutunian, and Y. Wang at the McGill Genome Centre for their services through the 10X platform, and M-H. Lacombe, E. Iourtchenko, and H. Pagé-Veillette at the Research Institute of the McGill University Health Centre Immunophenotyping Core for access to flow cytometry equipment and cell sorting services.

## Author contributions

T.A., S.L., C.A.P., and T.T. conceptualized the study and designed all the experiments. T.A., S.L., and L.A. conducted the experiments and T.A., S.L., and Z.L. curated the data and performed all the analyses. R.S., C.M., A.D., A.L., C.M., M.L., S.M., K.Y., S.M.S., C.A.P., and T.T. supplied resources and reagents. C.A.P. and T.T. obtained funding for this study. T.A., S.L., C.P., and T.T. wrote, reviewed, and edited the initial draft of the manuscripts. All authors reviewed and approved the submission of the manuscript.

## SUPPLEMENTARY MATERIALS

**Supplementary Table 1** – detailed patient characteristics (.xlsx)

**Supplementary Figure 1 –** Peripheral blood B cell profiling

**Supplementary Figure 2 –** Single-cell RNA-sequencing of B cells

**Supplementary Figure 3 –** B cell immunoglobulin gene usage

**Supplementary Figure 4 –** B cell clonal diversity

**Supplementary Figure 5 –** Clinical characteristics according to autoantibody seropositivity

**Supplementary Figure 6 –** Correlations between B cells and autoantibody titers

**Supplementary Figure 7 –** Correlations between plasmablasts and autoantibody titers

**Supplementary Figure 8 –** IgG deposition in kidneys of patients with idiopathic podocytopathies

**Supplementary Data File 1** – lists of differentially expressed genes for each B cell cluster against all other clusters (.csv)

**Supplementary Data File 2** – list of differentially expressed genes for total B cells in INS against controls (.csv)

**Supplementary Data File 3** – lists of differentially expressed genes for each B cell cluster in INS against controls (.xlsx)

---

**Supplementary Figure 1.**
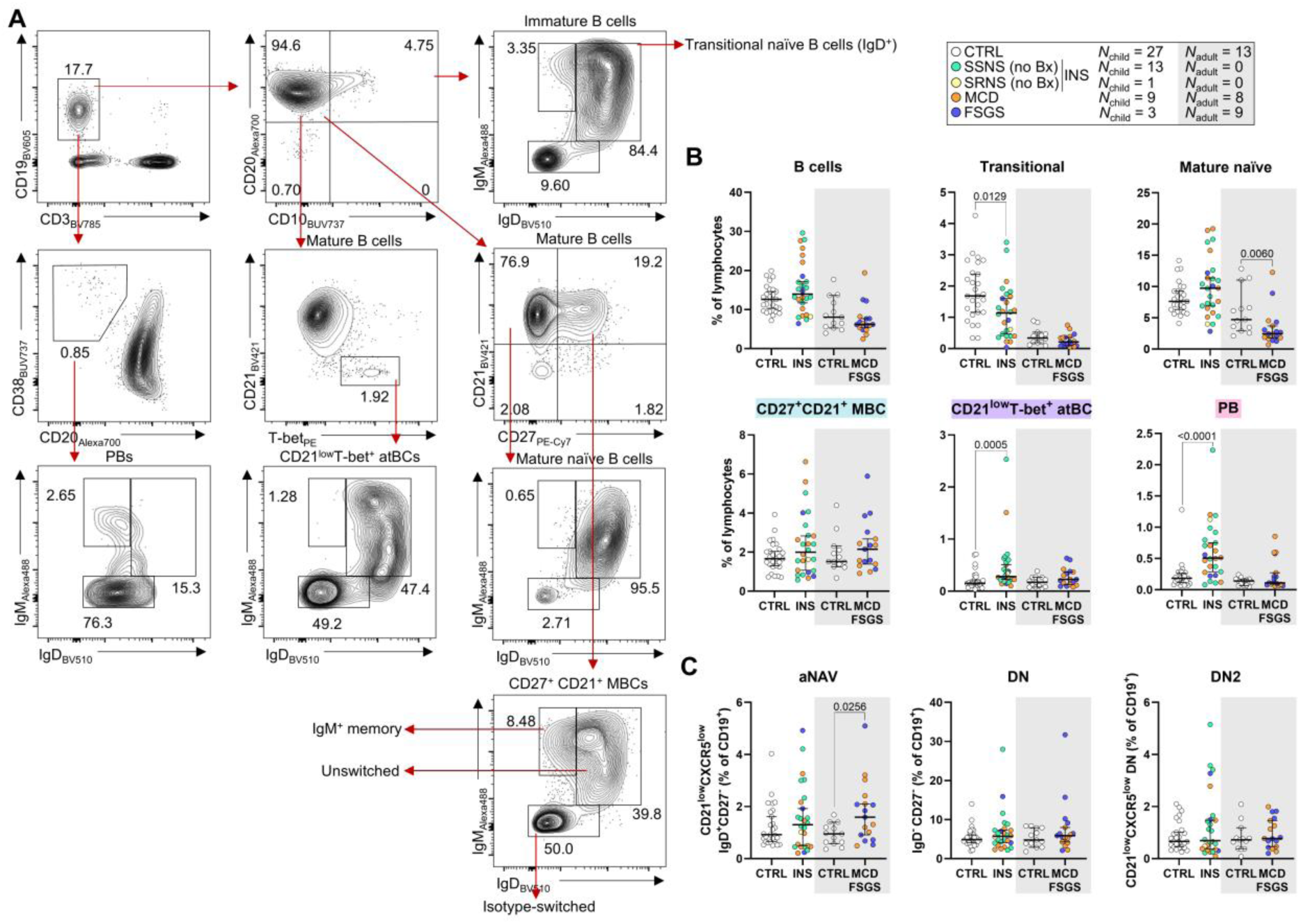
Peripheral blood B cell profiling. (**A**) Gating strategy for B cell populations. (**B**) Frequencies of total, transitional naïve, mature naïve, classical memory (MBC), and CD21^low^ T-bet^+^ atypical (atBCs) B cells and plasmablasts (PBs) within the lymphocyte pool. SSNS, steroid-sensitive nephrotic syndrome; SRNS, steroid-resistant nephrotic syndrome; MCD, minimal change disease; FSGS, focal segmental glomerulosclerosis. (**C**) Frequencies of activated naïve B (aNAV), double-negative B (DN), and DN2 B cells from total B cells. Each point refers to a single donor and summary statistics are expressed median with 95% confidence intervals. *P*-values were determined using Mann-Whitney *U*-tests comparing controls and patients in children and adults separately. This Supplementary Figure is associated with Figure 1.

**Supplementary Figure 2.**
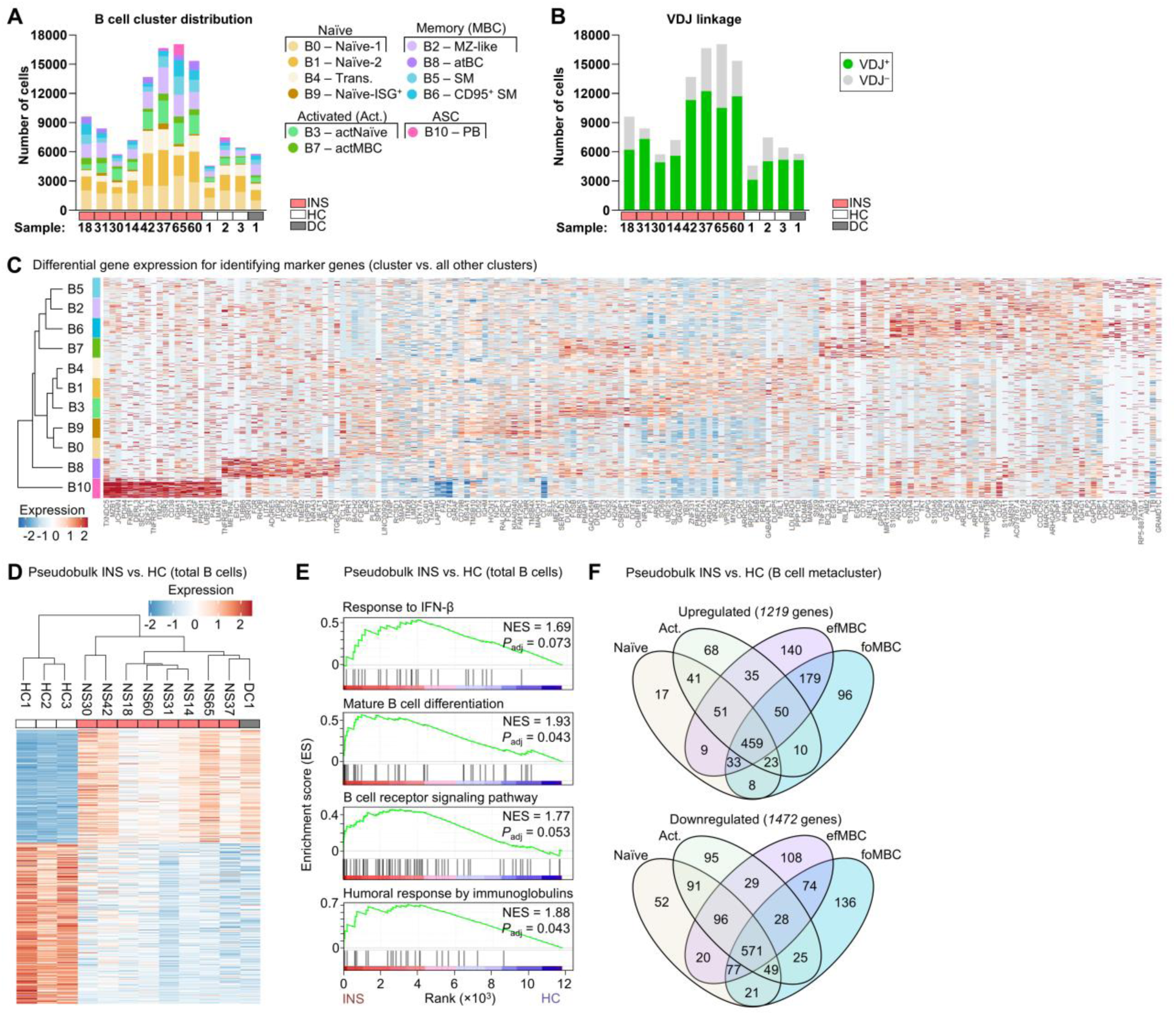
Single-cell RNA-sequencing of B cells. **(A)** Stacked bar graphs showing sample-level distribution of B cells from the single-cell RNA-sequencing dataset stratified by cluster. INS, idiopathic nephrotic syndrome; HC, healthy control; DC, disease control. **(B)** Stacked bar graphs showing sample-level distribution of B cells from the single-cell RNA-sequencing dataset stratified by having paired variable-diversity-joining (VDJ) sequences. **(C)** Heatmap of the top 20 expressed genes by each B cell cluster. **(D)** Hierarchical clustering of study participants and a heatmap of the differentially expressed genes between INS and healthy controls in total B cells. (**E**) Enrichment of gene ontology terms in total B cells from children with INS versus healthy controls. (**F**) Overlap of upregulated and downregulated differentially expressed genes in INS versus healthy controls for each B cell metacluster. Act., activated; efMBC, extrafollicular MBC; foMBC, follicular MBC. This Supplementary Figure is associated with Figure 2.

**Supplementary Figure 3.**
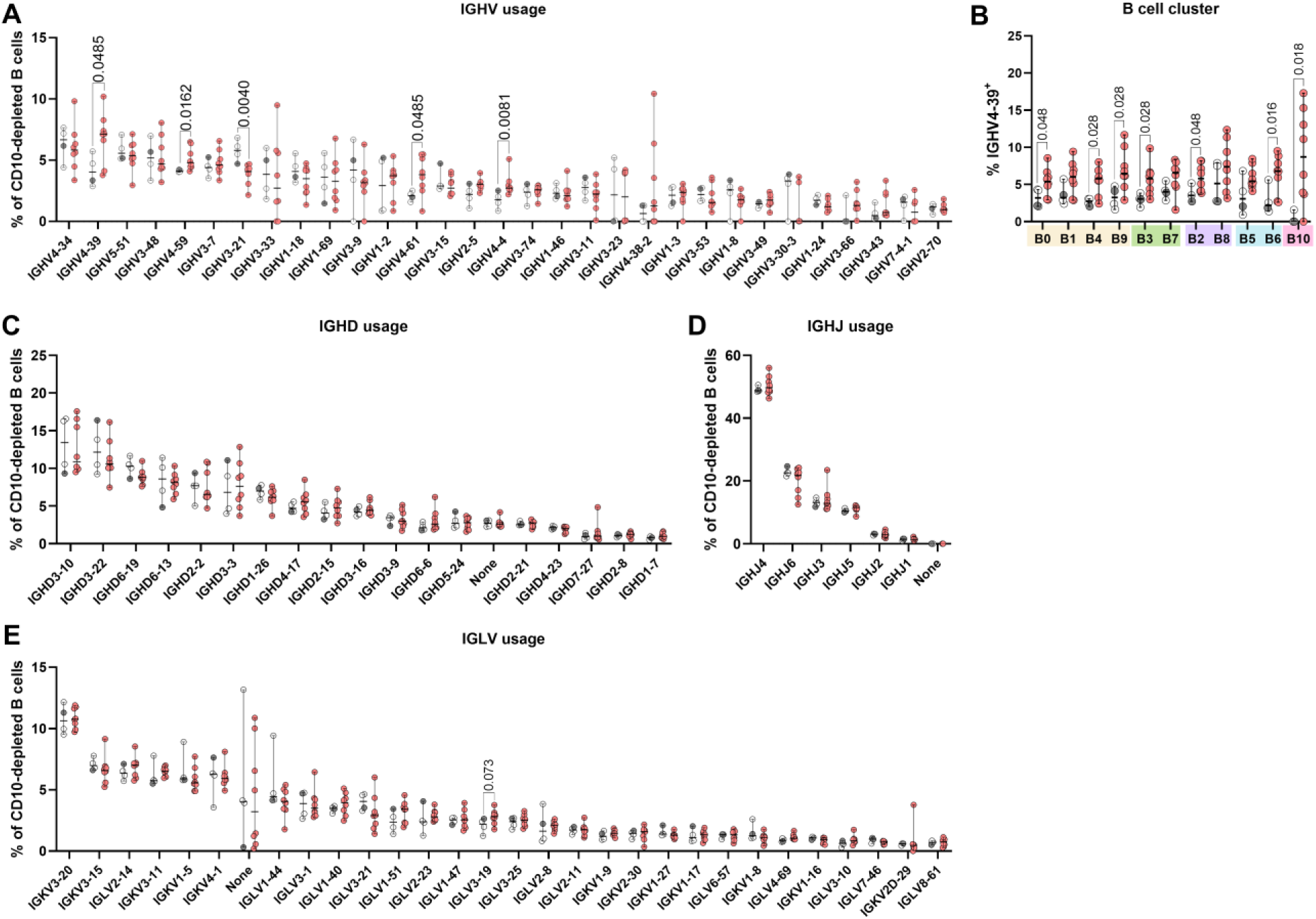
B cell immunoglobulin gene usage. (**A**) *IGHV* usage by total B cells. Only *IGHV* genes expressed by at least 1% of B cells are depicted. (**B**) Percentages of V_H_4-39^+^ cells within each B cell cluster. (**C-E**) *IGHD* (C), *IGHJ* (D), *IGLV* (E) usage by total B cells. Each dot represents a single patient with summary statistics represented as median with 95% confidence intervals. *P*-values were determined using independent Mann-Whitney *U*-tests between INS and controls. This Supplementary Figure is associated with Figure 3.

**Supplementary Figure 4.**
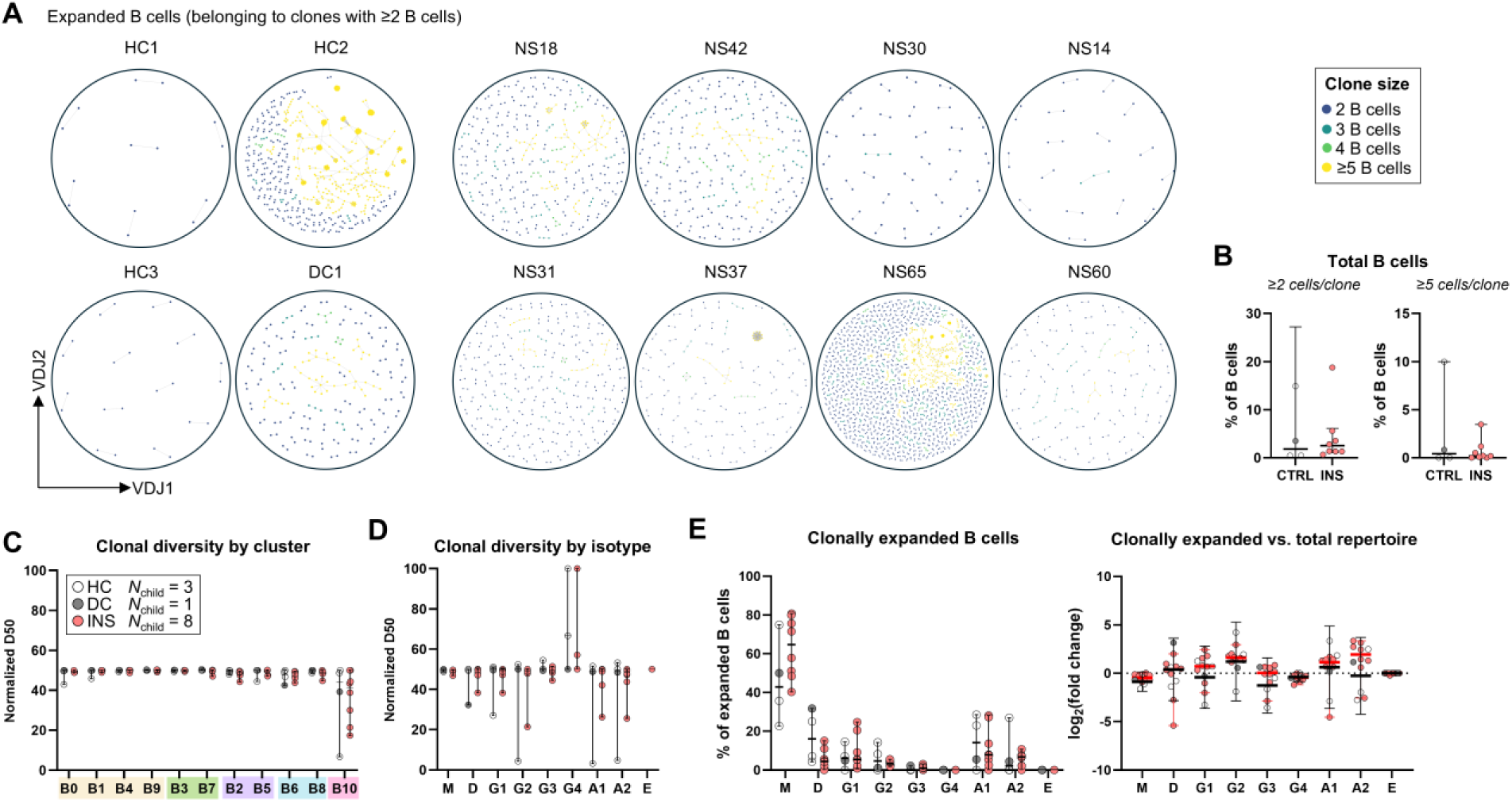
B cell clonal diversity. (**A**) Dandelion plots of clonally expanded B cells (at least two B cells within the clonotype) from each donor. (**B**) Frequencies of clonally expanded or highly expanded (at least five B cells within the clonotype) in children with INS or controls (CTRL). (**C-D**) Clonal diversity expressed using the normalized D50 index by B cell cluster (C) and isotype (D). (**E**) Isotype frequencies within the clonally expanded pool (*left*) and their log_2_(fold change) from isotype frequencies within the total B cell pool. Each dot represents a single donor with summary statistics represented as median with 95% confidence intervals. *P*-values were determined using independent Mann-Whitney *U*-tests between INS and controls. This Supplementary Figure is associated with Figure 3.

**Supplementary Figure 5.**
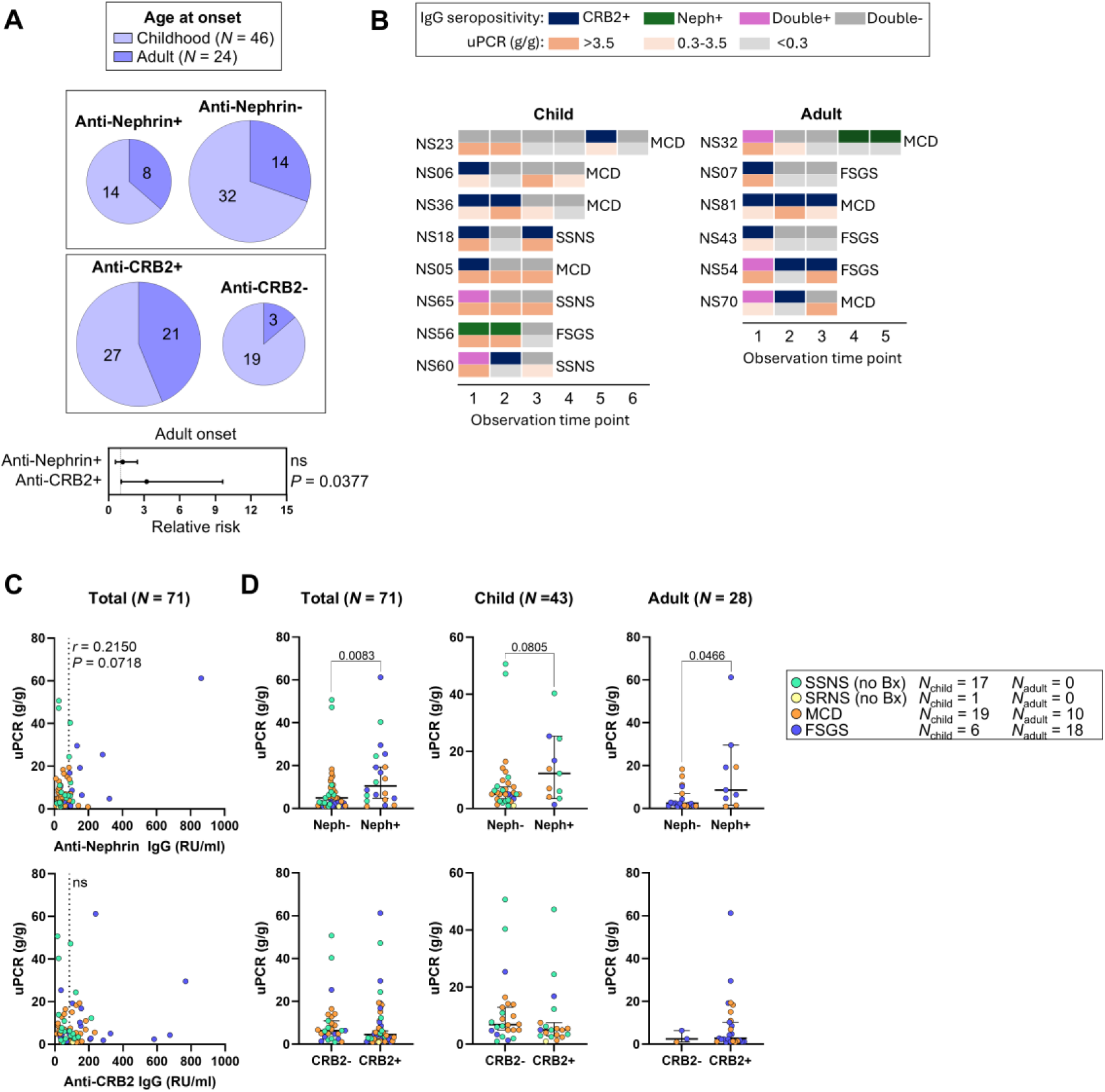
Clinical characteristics according to autoantibody seropositivity. (**A**) Numbers of patients with disease onset in childhood or adulthood stratified by autoantibody seropositivity (*top*). The size of each pie chart is proportional to the number of individuals captured in that group. Forest plots of the relative risk for adulthood onset given anti-CRB2 or anti-Nephrin seropositivity (*below*). **(B)** Timeline of autoantibody seropositivity status and proteinuria severity expressed as the protein-to-creatinine ratio (uPCR) in patients with samples available at three or more independent clinical visits. **(C)** Correlation of uPCR and IgG autoantibody titers in patients with INS (*N*_child_ = 18), including SSNS (*N*_child_ = 17) and SRNS (*N*_child_ = 1), or MCD (*N*_child_ = 19, *N*_adult_ = 10) and primary FSGS (*N*_child_ = 6, *N*_adult_ = 18). **(D)** uPCR graphed by autoantibody seropositivity for the total cohort or for children and adults independently. In C and D, each dot represents a single donor and summary statistics are shown as median with 95% confidence intervals. *P*-values were determined using Fisher’s exact tests in A and Mann-Whitney *U*-tests in C and D. This Supplementary Figure is associated with Figure 4.

**Supplementary Figure 6.**
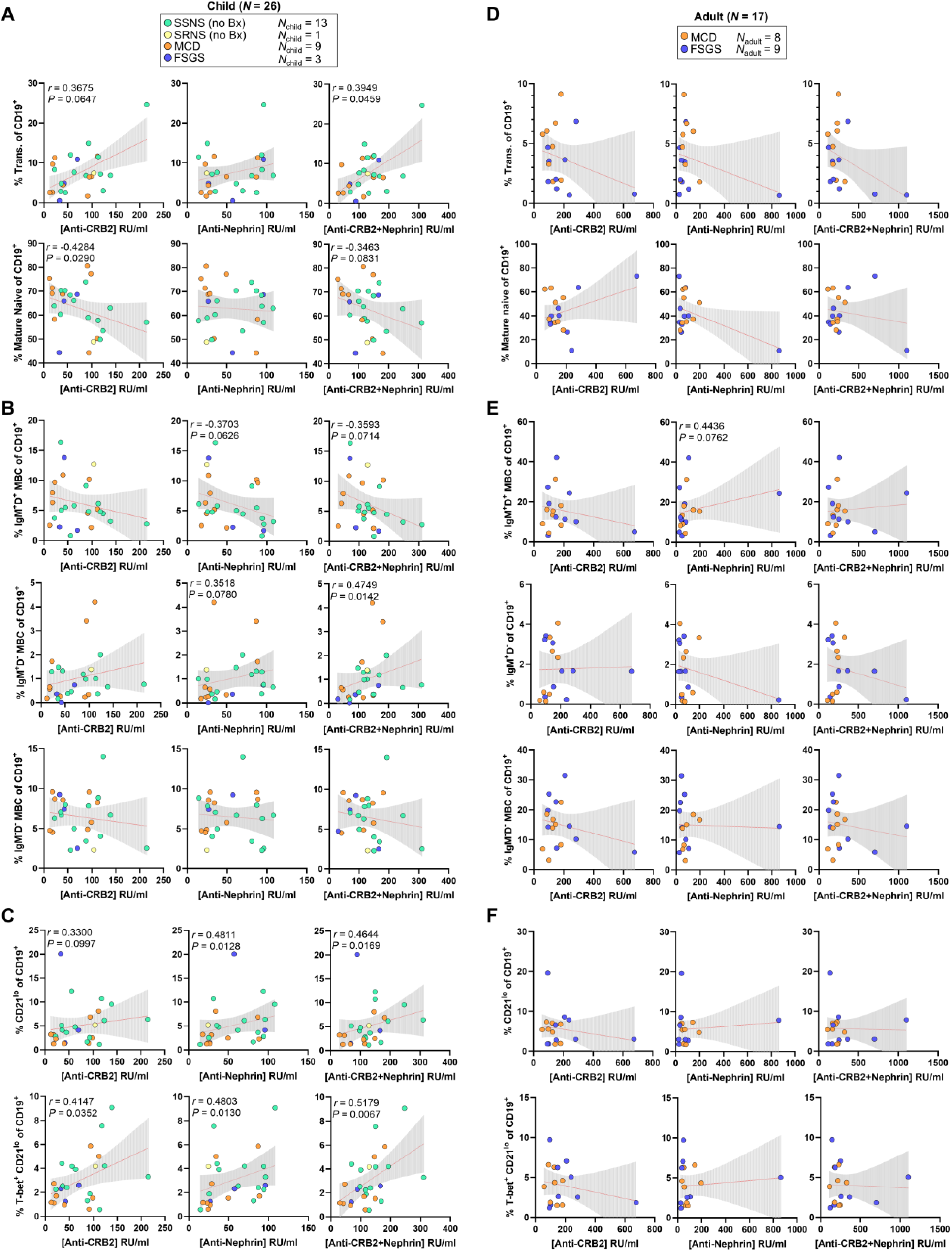
Correlations between B cells and autoantibody titers. **(A)** Correlation between the frequency of transitional B cells or mature naïve B cells with serum titers of anti-Nephrin IgG, anti-CRB2 IgG or their combined titers in children. **(B)** Correlation between the frequency of unswitched (IgM^+^ IgD^+^), IgM (IgM^+^ IgD^-^), and isotype-switched (IgM^-^ IgD^-^) MBC B cells with serum titers of anti-Nephrin IgG, anti-CRB2 IgG or their combined titers in children. **(C)** Correlation between the frequency of CD21^low^ and T-bet^+^ CD21^low^ B cells with serum titers of anti-Nephrin IgG, anti-CRB2 IgG or their combined titers in children. **(D)** Correlation between the frequency of transitional B cells or mature naïve B cells with serum titers of anti-Nephrin IgG, anti-CRB2 IgG or their combined titers in adults. **(E)** Correlation between the frequency of unswitched, IgM, and isotype-switched MBC B cells with serum titers of anti-Nephrin IgG, anti-CRB2 IgG or their combined titers in adults. **(F)** Correlation between the frequency of CD21^low^ and T-bet^+^ CD21^low^ B cells with serum titers of anti-Nephrin IgG, anti-CRB2 IgG or their combined titers in adults. Each dot represents a single donor and simple linear regression lines with 95% confidence intervals are provided for visualization. Non-parametric Spearman correlations were performed to obtain *r*- and *P*-values. This Supplementary Figure is associated with Figure 4.

**Supplementary Figure 7.**
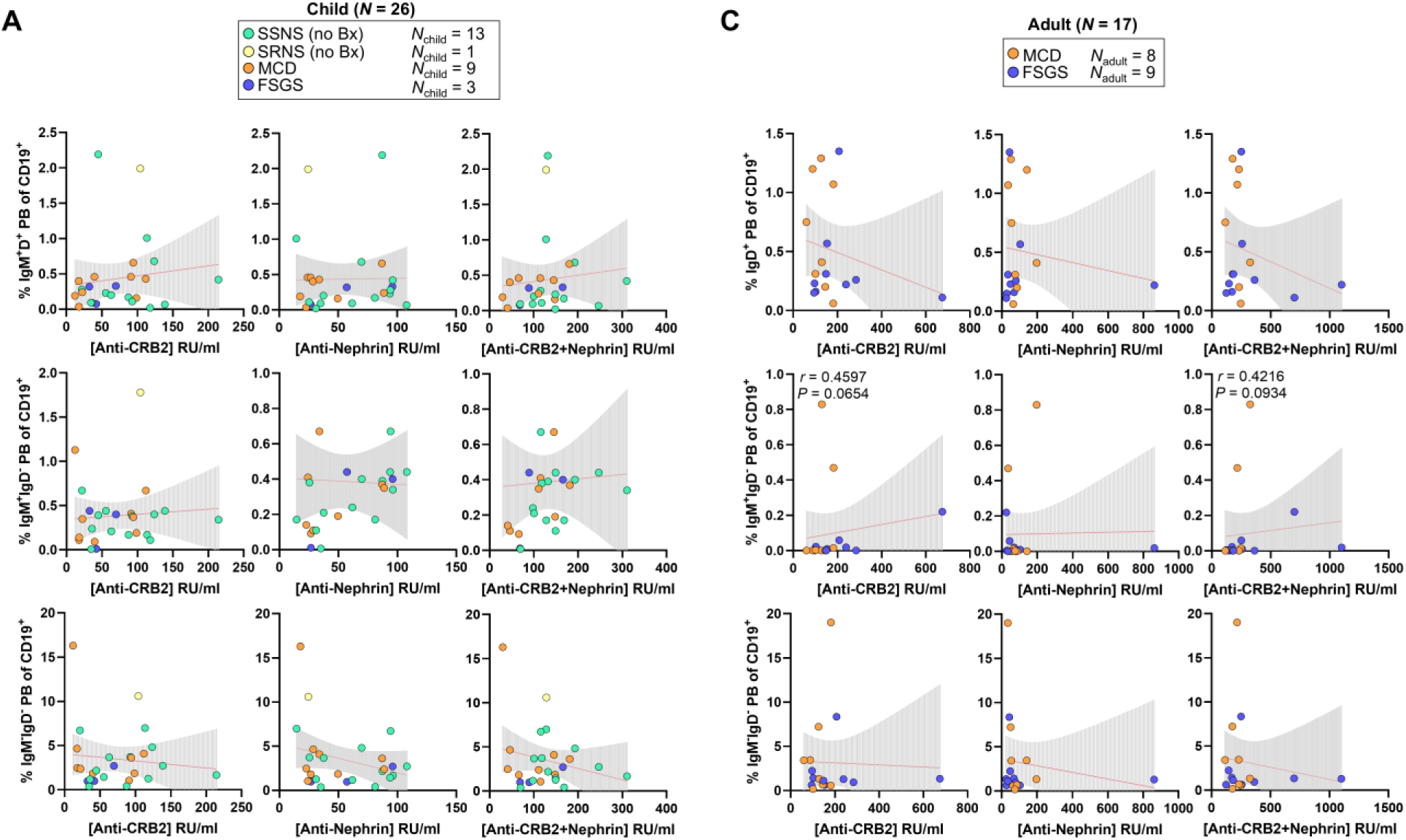
Correlations between plasmablasts and autoantibody titers. (**A-B**) Correlation between the frequency of unswitched (IgM^+^ IgD^+^), IgM (IgM^+^ IgD^-^) and isotype-switched (IgM^-^ IgD^-^) plasmablasts with serum titers of anti-Nephrin IgG, anti-CRB2 IgG or their combined titers in children (A) and adults (B). Each dot represents a single donor and simple linear regression lines with 95% confidence intervals are provided for visualization. Non-parametric Spearman correlations were performed to obtain *r*- and *P*-values. This Supplementary Figure is associated with Figure 4.

**Supplementary Figure 8.**
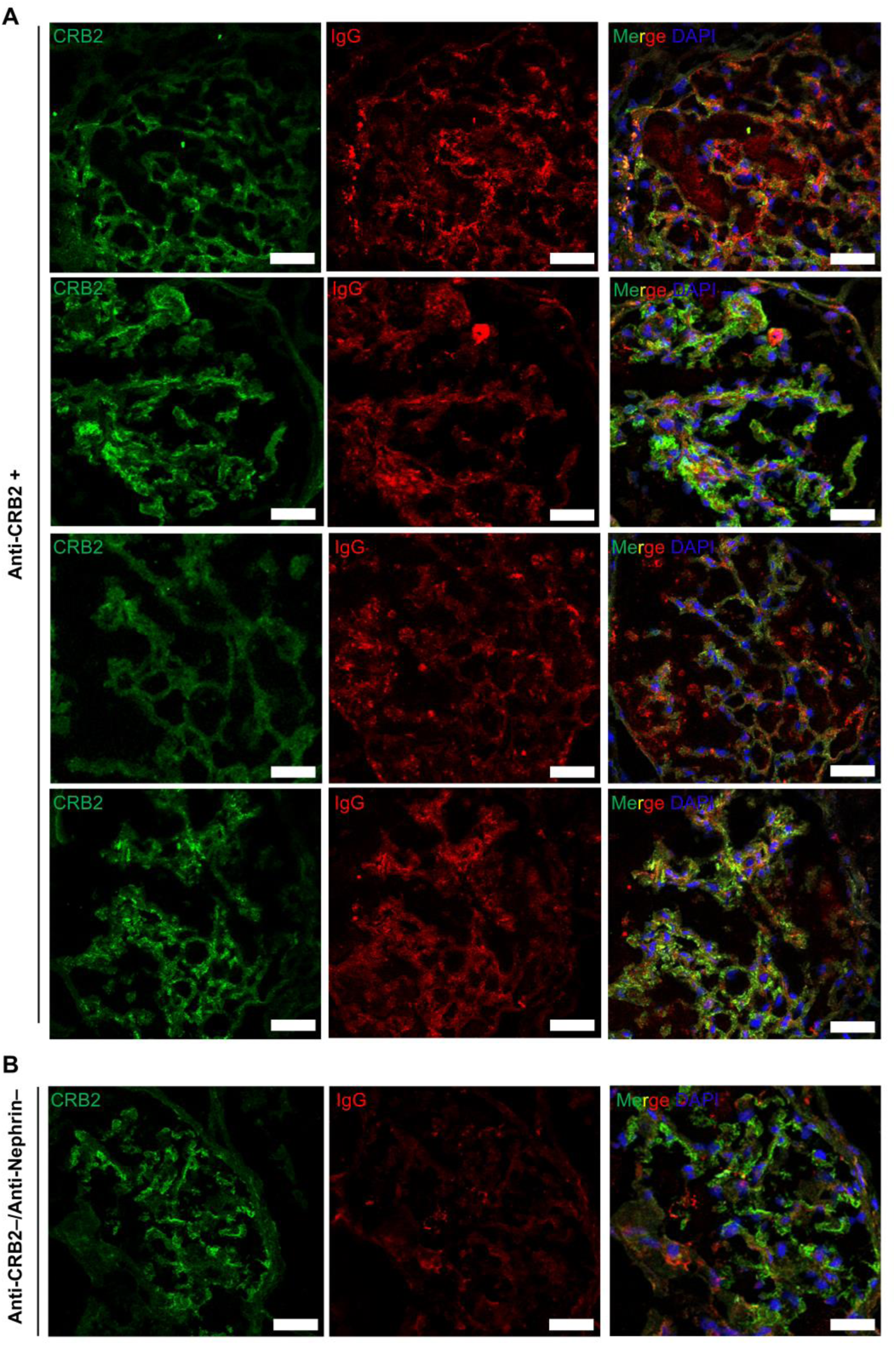
IgG deposition in kidneys of patients with idiopathic podocytopathies. (**A-B**) Representative glomeruli from flash frozen kidney biopsies from the remaining four anti-CRB2 IgG seropositive patients (A) and one seronegative patient (B) present in the quantification in Figure 5B showing CRB2 (*red*) and IgG (*red*) co-localization. This Supplementary Figure is associated with Figure 5.

**Supplementary Table 1. Patient characteristics.** This table lists all the patients included in the study and their characteristics. The manuscript code is the anonymized ID of each patient included in this manuscript. The diagnosis column represents the disease with which the patient was diagnosed, where INS denotes unbiopsied idiopathic nephrotic syndrome, SSNS = steroid-sensitive INS, SRNS = steroid-resistant INS, MCD = minimal change disease, FSGS = primary focal segmental glomerulosclerosis, MN = membranous nephropathy, IgAN = IgA nephropathy, GEN = genetic disease, C3GN = C3 glomerulonephritis, FIB = fibrillary glomerulonephritis, A-GBM = anti-glomerular basement membrane disease, AHUS = atypical hemolytic uremic syndrome, MGRS = monoclonal gammopathy of renal significance, PGNMID = proliferative glomerulonephritis with monoclonal immune deposits, and SLE = systemic lupus erythematosus. Steroid responsiveness denotes if patients were steroid-sensitive (SSNS) or resistant (SRNS). Sample date represents the date of collection of the first tested sample for each patient. The age column denotes the age of each patient at the time of sample collection. For the sex, F = female and M = male. The nest three columns describe the treatments received by each patient: if they were receiving steroids at the time of sample collection and the treatment duration, if they were receiving any other immunosuppressive agents at the time of sample collection (MMF = mycophenolate mofetil, RTX = rituximab), and if they were previously exposed to RTX before sample collection. Disease status includes active disease, defined as proteinuria >1 g/day and/or a diagnosis of relapse by the attending nephrologist, and remission, defined as proteinuria <0.3 g/day and/or diagnosis of remission by the nephrologist (more details are provided in the Methods section). uPCR denotes urinary protein-to-creatinine ratio, measured here in g/g. The next four columns state the anti-CRB2 and the anti-Nephrin autoantibody titer in µg/mL and the associated interpretation (seropositivity or seronegativity). The final columns titled IgM/IgA isotyping, Confocal, Super resolution, Flow cytometry, scRNAseq (single-cell RNA-sequencing), and *in vitro* plasmablast differentiation indicates the patients that were included in each of these analyses.

